# Pharmacokinetic Acceleration via CYP3A4 Hyperactivation as a Clinically Actionable Mechanism of Targeted Therapy Resistance in NSCLC

**DOI:** 10.64898/2026.05.16.725647

**Authors:** Pragya Kumar, Menkara Henry, Julia Pelesko, Bina Desai, Malgorzata Tyczynska Weh, Ramu Kakumanu, Min Liu, Natalia Souza Nunes Siqueira, Robert Vander Velde, Viktoriya Marusyk, Jyllica Aurelio, Eric Haura, Jhanelle Gray, Bruna Pellini, David Basanta, Andriy Marusyk

## Abstract

Resistance of cancers to targeted therapies is traditionally framed as a tumor-intrinsic phenomenon, mediated by tumor cell-intrinsic or microenvironmental mechanisms. Here, we identify a tumor-extrinsic, systemic resistance mechanism resulting from hyperactivation of the hepatic cytochrome P450 enzyme, CYP3A4. This tumor-extrinsic resistance mechanism can function independently of, or in tandem with, tumor-intrinsic resistance. Focusing on experimental mouse models of targetable lung cancer, we find that xenobiotic-mediated induction of CYP3A4 results in accelerated drug metabolism and a drastic reduction in systemic and tumor-drug exposure *in vivo*. CYP3A4 activation can be triggered by chemically unrelated xenobiotics, leading to resistance to a wide range of targeted therapies, including ALK, EGFR, and KRASG12C inhibitors. Retrospective analysis of clinical cohorts suggests that variability in CYP3A4 activity might be a major contributor to variability in clinical outcomes. While higher CYP3A4 activity leads to sub-therapeutic tumor drug exposure and shorter progression-free survival, reduced drug metabolism is expected to result in supratherapeutic exposure and increased systemic toxicity. To address the consequences of abnormal CYP3A4 activity, we utilized mathematical modeling to demonstrate that drug concentrations can be restored through the optimization of dosing amounts and intervals. Further, we show that tumor sensitivity to targeted therapies can be rescued through pharmacological inhibition of CYP3A4. Our findings establish systemic metabolic variability as a *bona fide* resistance and toxicity driver, providing a translational framework for personalized dosing to maximize both safety and efficacy.

## Introduction

Targeted therapies that suppress oncogenic signaling addictions can induce strong and often durable clinical responses with relatively low systemic toxicities, making them desirable frontline therapy choices. Unfortunately, a subset of cancers fails to respond (intrinsic resistance), or show weak, transient responses, despite the presence of a strong oncogenic driver. Moreover, tumors that show strong initial responses eventually develop resistance (acquired resistance) and relapse^1–4^. Both intrinsic and acquired resistance are generally attributed to tumor-intrinsic mechanisms. Historically, consideration of resistance mechanisms has been focused on mutational mechanisms that impair drug binding, change drug-target stoichiometry via drug amplification, or activate bypass signaling by activation of alternative signaling genes^2,4^. Subsequently, consideration of resistance mechanisms has been extended to epigenetic mechanisms, such as transcriptional upregulation of bypass signaling pathways, enhanced stemness, and lineage transition^3,5–7^. Finally, environment-mediated resistance has recently emerged as a major contributor to therapeutic failures via paracrine interactions between tumor cells and various cellular components of the tumor microenvironment, mediated by metabolites, exosomes, and growth factors, as well as juxtacrine interactions between tumors and some components of the extracellular matrix^8–10^. In all these various tumor-intrinsic mechanisms, resistance results from the reduction or complete loss of sensitivity of tumor cells to the cytostatic and cytotoxic effects of therapeutic agents.

While trying to identify collateral sensitivities of tumor cells capable of survival under ALK inhibitors (ALKi) in non-small cell lung cancers (NSCLC), we have encountered a puzzling discrepancy between the *in vitro* and *in vivo* effects of a KDM4 inhibitor, JIB-04. While JIB-04 co-treatment enabled the elimination of ALKi persisters *in vitro*, it led to ALKi resistance *in vivo*. We found that this discrepancy reflected the JIB-04-mediated upregulation of CYP3A4, a P450 family member responsible for the metabolic breakdown and accelerated excretion of multiple therapeutic agents^11^. CYP3A4 activation accelerated drug pharmacokinetics (PK), leading to a quick drop of plasma and tumor drug concentrations, resulting in lower tumor drug exposure. Resistance-promoting effects of JIB-04 can be phenocopied by chemically unrelated CYP3A4 activators. CYP3A4 activation promoted resistance to multiple additional targeted therapies, including clinically relevant EGFR and KRAS inhibitors, suggesting a broad generalizability of our findings. Whereas JIB-04 is not a clinically relevant drug, multiple dietary, pharmacological, and genetic factors can modulate CYP3A4 activity in patients, which can result in >10x patient-patient variability in this enzyme^12^. Our pilot analyses indicate that enhanced CYP3A4 levels might be a major cause of clinical therapy resistance. Using integration of mathematical and experimental modeling, we demonstrate that this resistance mechanism can be mitigated through clinically actionable therapeutic strategies involving CYP3A4 inhibitors or altered dosing/scheduling regimens.

## Results

### Despite sensitizing ALK+ NSCLC cells to ALKi *in vitro, JIB04* causes resistance to ALKi *in vivo*

Upon exposure to targeted therapies, a subset of tumor cells is typically capable of surviving elimination (persistence) at the expense of reduced proliferation rates. Over time, persisters adapt to therapeutic selection pressures, eventually evolving resistance (net positive growth under treatment). The transition from persistence to resistance often involves substantial changes in gene expression networks that can be conceptualized as a resistance continuum^5,13^. Subsequently, evolving populations can be exquisitely sensitive to pharmacological inhibitors of epigenetic machinery, involved in rewiring gene expression networks^7,14^. Consistent with the previous findings in EGFR mutant NSCLC^14^, we found that ALK+ NSCLC cells persisting under clinically relevant concentrations of ALKi alectinib, lorlatinib and crizotinib, are much more sensitive to a histone demethylase inhibitor, JIB-04 compared to therapy naïve or ALKi resistant cells **(Fig. S1A)**. Subsequently, JIB-04 strongly augments responses of ALK+ cell lines H3122 and STE1 to ALKi, leading to a complete elimination of tumor cells in 3D *in vitro* cultures **(Fig. 1A-C)**.

**Figure 1:**
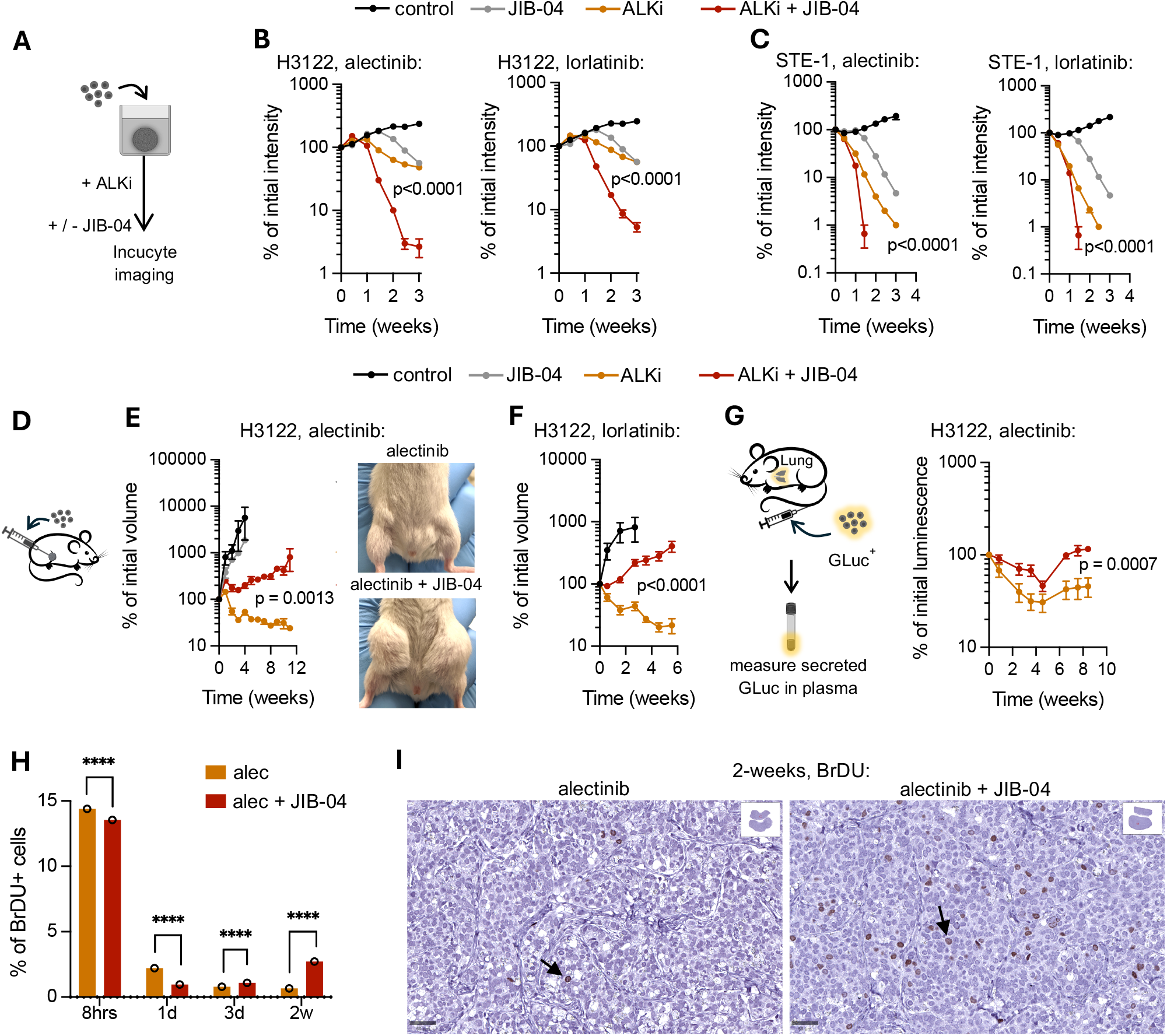
JIB-04 causes resistance to ALKi *in vivo*, despite improving sensitivity *in vitro*. **A**. Schemata of *in vitro* sensitivity assay of ALK+ cells for panels B, C. Cells were grown as 3D spheroids in low-attachment plates and treated as indicated. Fluorescence intensity was measured for quantitation. **B, C**. Sensitivity of cells to indicated treatments (0.5mM alectinib (alec) and 2mM lorlatinib (lor) with or without 2.5mM JIB-04) with DMSO as vehicle in long-term sensitivity assay using spheroids of H3122 cells or STE-1 cells. Intensity was normalized as % of pre-treatment intensity. Data represents n = 3 replicates for all groups and 2-way ANOVA with Sidak’s test was performed (p-value denotes interaction). **D**. Schemata showing model of subcutaneous xenografts. Cells were injected subcutaneously in the flanks of NSG mice for panels E, F. **E**. Change in tumor volume of H3122 subcutaneous xenografts over time under indicated treatments. Y axis shows the % volume, normalized with initial volume measurements at the start of treatment. n = 9 tumors for control and alec+JIB-04, n = 6 tumors for JIB-04 and n = 10 tumors for alec alone. Right panel shows representative images of the tumors with indicated treatments. p-value represents interaction term of 2-way ANOVA with Sidak’s test. **F**. Change in tumor volume of H3122 subcutaneous xenografts over time under indicated treatments. Y axis shows the % volume, normalized with initial volume measurements at the start of treatment. n = 4 tumors for control and n = 10 tumors for lor and lor+JIB-04. p-value represents interaction term of 2-way ANOVA with Sidak’s test. **G**. Schemata showing model of orthotopic injection of H3122 cells tagged with Gaussia secreted luciferase plasmid into tail vein. Luminescence was measured in blood-plasma every week and normalized to luminescence at the start of indicated treatments (shown in y-axis). n = 5 mice for both groups. 2-way ANOVA with Sidak’s test was used for all data in the figure and p-value denotes interaction. **H**. BrDU staining on H3122 xenograft tumors harvested after indicated timepoints. % BrDU cells are calculated relative to total number of cells analyzed for indicated treatments. Chi-square test was performed using cumulative number of positive and negative cells counted (total ∼40,000-230,000 cells across groups) and p-value is shown. **I**. Histology images (scalebar at 50mm, scanned at 20X) are shown for the 2-week timepoint. Arrows point to BrDU+ cells. For all *in vivo* experiments, dosages used were, 25mg/kg alectinib, 10mg/kg lorlatinib and 55mg/kg JIB-04. Mean ± SEM are shown for all graphs, *p < 0.05, **p < 0.01, ***p < 0.001, and ****p < 0.0001.

Surprisingly, when we attempted to validate the effects of the ALKi-JIB-04 combination *in vivo*, the opposite effect was observed. As expected^13^, alectinib and lorlatinib, administered as a single agent, induced strong and durable regression of subcutaneous H3122 xenograft tumors **(Fig. 1D-F)**. In contrast, co-treatment with JIB-04 not only failed to enhance the responses to ALKi but also led to a rapid onset of resistance **(Fig. 1E, F)**. To assess whether the same unexpected reversal occurs in an orthotopic context, we used tail vein injections to seed H3122 cells into the lungs. The cells were engineered to express Gaussia luciferase (GLuc), enabling quantitative monitoring of tumor burden from *ex vivo* analyses of blood plasma^15^ **(Fig. 1G)**. Similar to subcutaneous tumors, JIB-04 co-treatment strongly desensitized lung H3122 tumors to alectinib **(Fig. 1G)**. The paradoxical reversal of the effect of JIB-04 between *in vitro* and *in vivo* was also observed in a different model of ALK+ NSCLC, the STE1 cell line **(Fig. S1B)**.

Given the slight delay in the full onset of desensitization in xenograft tumor models, we examined the impact of JIB-04 on *in vivo* proliferation dynamics using the BrdU label, which marks cells in the S phase of the cell cycle. Notably, the alectinib-JIB-04 combination resulted in strong initial suppression of the proliferation of H3122 tumor cells that was similar to or stronger than the suppression in mice treated with alectinib monotherapy **(Fig. 1H, S1C, D)**. However, by 2 weeks, cell proliferation was partially restored under the alectinib-JIB-04 combination **(Fig. 1H, I, S1D)**, consistent with the resumption of partially repressed but net-positive tumor growth.

### *In vivo* resistance is linked with enhanced systemic drug clearance

The striking discrepancy between the *in vitro* and *in vivo* effects of JIB-04 indicated a modulating impact of either systemic or microenvironmental context. A recent clinical study indicated that patients treated with identical doses of alectinib display strong variability in plasma drug concentrations and that lower drug concentrations correlate with worse clinical outcomes^16^. Therefore, we asked whether JIB-04 impacts systemic levels of ALKi. Mass spectrometry-based analyses revealed a drastic reduction in alectinib and lorlatinib levels in blood and tumor tissues of mice co-treated with JIB-04, compared to the ALKi monotherapies **(Fig. 2A, S2A)**, implicating reduced ALKi exposure of tumor cells as a likely cause of resistance.

**Figure 2:**
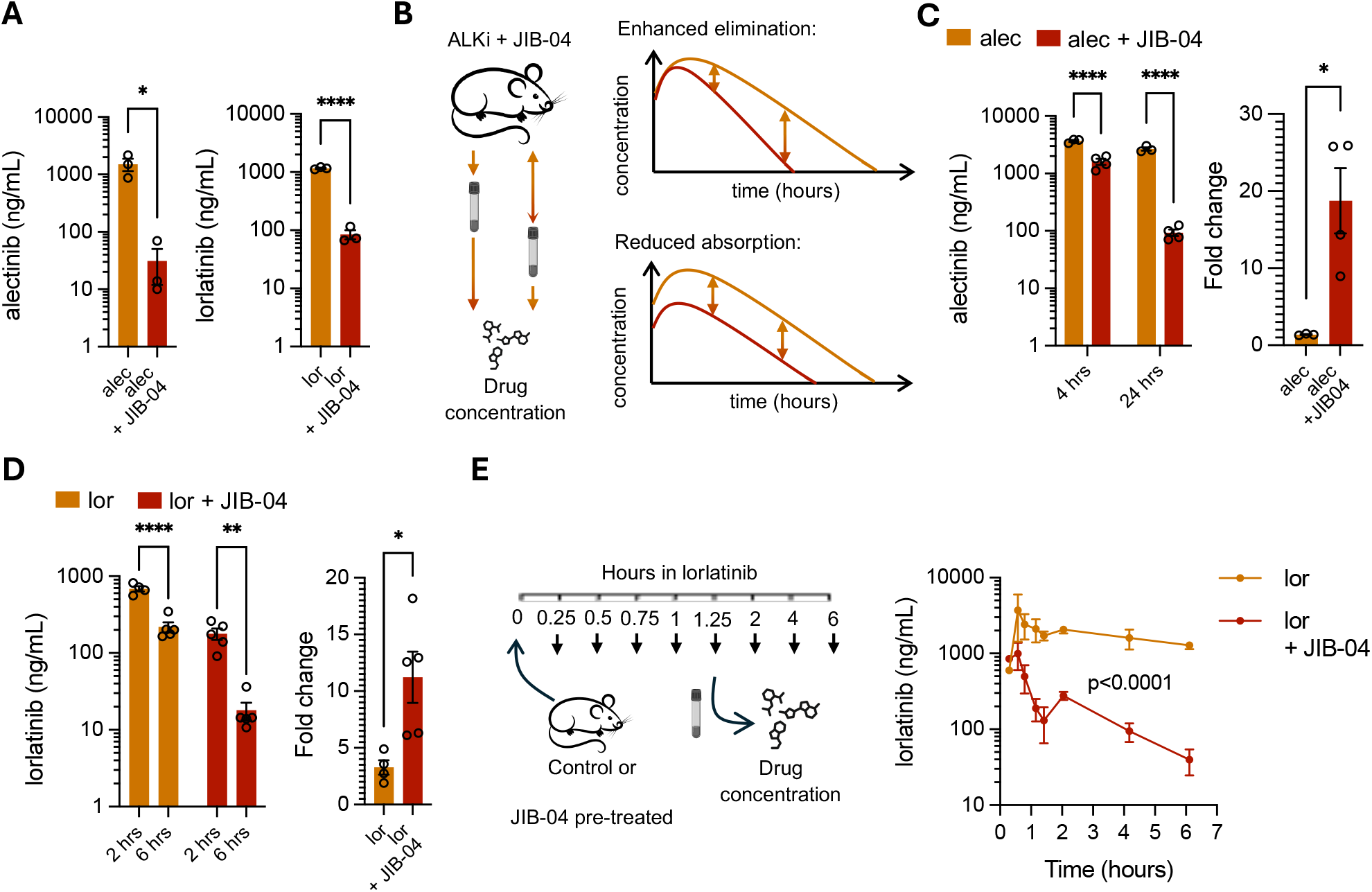
JIB-04 accelerates ALKi elimination. **A**. Concentrations in plasma of alectinib and lorlatinib from mice bearing H3122 tumors and treated with alectinib for 2.5 weeks or lorlatinib for 3.7 weeks and JIB-04 respectively as indicated. n = 3 for all groups. Unpaired t-test was performed. **B**. Schemata showing experimental design for panels C and D and expected drug concentration dynamics. Mice were treated with indicated drugs and blood was collected at endpoint at 2 timepoints after last dose of ALKi. Enhanced elimination shows larger difference in concentrations at late vs early timepoint while reduced absorption shows the same variation. **C**. Left: plasma concentrations of alectinib from mice pre-treated with JIB-04 for 10 days. Blood was collected at indicated hours after the single dose of alectinib. p-value represents interaction of 2-way ANOVA with Sidak’s test. Right: fold change of concentrations at 24 hours relative to 4 hours. n = 3 mice for alec and n = 4 mice for alec+JIB-04. Welch t-test was performed. **D**. Left: plasma concentrations of lorlatinib from mice treated with indicated drugs for 5.5 weeks. Blood was collected at indicated hours. p-value represents interaction of 2-way ANOVA with Sidak’s test. Right: fold change of concentrations at 6 hours relative to 2 hours. n = 5 mice for 6-hour timepoint, n = 4 mice for lor 2 hours and n = 5 mice for lor+JIB-04 2 hours timepoint. Welch t-test was performed. **E**. Schemata showing experimental design and pharmacokinetic analysis of lor in control or JIB-04 pre-treated mice (for 10 days). Blood-plasma was collected at indicated hours after single dose of 10mg/kg lor. n = 3 mice per timepoint for both groups. Mean ± SEM are shown. 2-way ANOVA with Sidak’s test was performed. Mean ± SEM are shown for all graphs, *p < 0.05, **p < 0.01, ***p < 0.001, and ****p < 0.0001.

The reduction in systemic levels of small-molecule oral drugs like ALKi could potentially reflect either reduced absorption or enhanced elimination (through metabolism and excretion)^17,18^. We reasoned that distinction between these two possibilities can be achieved by examining blood drug concentrations at distinct time points after a single dose of alectinib in control or JIB-04 pre-treated mice **(Fig. 2B)**. While alectinib blood concentrations were lower in JIB-04-treated mice at both 4- and 24-hour time points, the difference was much more pronounced at 24 hours in the JIB-04 pre-treated group, as the drug levels dropped more strongly **(Fig. 2C)**. Similar results were observed for lorlatinib **(Fig. 2D)**; more closely spaced time points were chosen to account for the higher lorlatinib PK velocity in mice^19,20^. These results indicate that the effect of JIB04 is attributable to the enhanced drug elimination. To validate this inference, we measured blood concentrations of lorlatinib with a single dose of lorlatinib in control and JIB-04 pre-treated mice at a higher temporal resolution **(Fig. 2E)**. Consistent with the expectations, we found a significant reduction in the concentration of lorlatinib in JIB-04 pre-treated mice versus control mice, particularly in the elimination phase **(Fig. 2E)**. JIB-04 co-treatment led to a ∼7-fold reduction in the plasma drug exposure for the first six hours following drug administration (area under the curve, AUC_0-6_), supporting our inference that the reduction of systemic ALKi levels in JIB-04-treated mice reflects enhanced drug elimination.

### Resistance-inducing PK acceleration is mediated by enhanced CYP3A11 expression

Elimination of small molecule drugs is primarily mediated by P450 family liver enzymes that metabolically convert xenobiotics, leading to their inactivation and/or enhanced excretion^17,21,22^. Thus, we examined the impact of JIB-04 on liver tissue transcriptomes. We found 197 genes significantly (p-value<0.05; with over 1.5 log2 fold change) altered between the control and JIB-04 treated groups **(Fig. S2B)**. Pathway enrichment analysis (KEGG, Reactome) of these differentially expressed genes revealed alterations in multiple pathways, including those involved in the inactivation of xenobiotics **(Fig. 3A and S2C, D)**. JIB-04 treatment enhanced the expression of multiple metabolic genes, including P450 (CYP) family member CYP3A11 **(Figure S2E)**. The murine CYP3A11 gene is an orthologue of the human CYP3A4 gene, which is the main mediator of metabolic conversion of oral small-molecule drugs, including ALKi, alectinib, and lorlatinib^23–25^. Transcriptional upregulation of CYP3A11 by JIB-04 was associated with the elevated protein expression in murine livers; this effect was independent of the ALKi co-treatment **(Fig. 3B)**.

**Figure 3:**
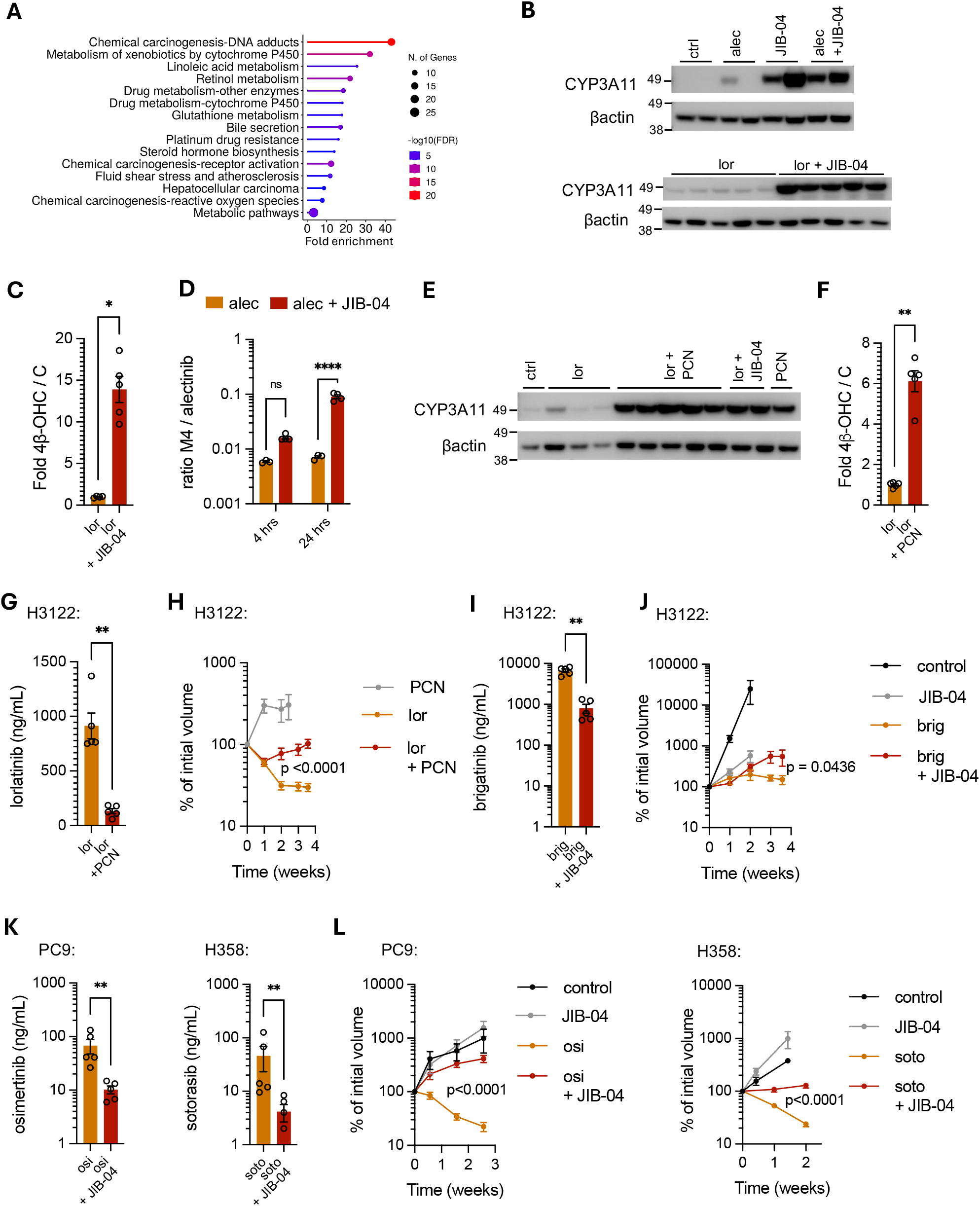
CYP3A11 hyperactivation leads to faster substrate drug elimination and resistance. **A**. Top 15 KEGG pathways using ShinyGO. Genes used have 1.5<log2 fold change in JIB-04 vs control, derived from differential gene expression analysis (p-value<0.05). **B**. Immunoblotting images showing protein levels of CYP3A11 with βActin as loading control. Proteins were harvested from mice treated with indicated drugs for 1.4 weeks for alec experiment and 5.5 weeks for lor experiment. **C**. Fold of ratio of plasma concentrations of 4β-hydroxy cholesterol with cholesterol (4β-OHC/C) where ratio in lor+JIB-04 group is normalized with ratio in lor alone group. Blood was collected from mice treated with indicated drugs for 5.5 weeks. n = 4 mice for lor and n = 5 mice for lor+JIB-04. Mann-Whitney test was performed. **D**. Ratio of concentrations of M4 over alec in plasma are shown. Mice were pre-treated with JIB-04 for 10 days followed by a single dose of alectinib and samples were collected at indicated timepoints. n = 3 mice for alec and n = 4 mice for alec+JIB-04. p-value represents interaction of 2-way ANOVA with Sidak’s test. **E**. Western blot images showing protein levels of CYP3A11 with βActin as loading control. Proteins were harvested from mice treated with indicated drugs for 3.5 weeks. **F**. Fold of 4β-OHC/C ratio where ratio in lor+JIB-04 group is normalized with ratio in lor alone group. Blood was collected from mice treated with indicated drugs for 3.5 weeks. n = 5 mice for all groups. **G**. Plasma concentrations of lorlatinib from mice treated with indicated drugs for 3.5 weeks. n = 5 mice for all groups. **H**. Change in tumor volume of H3122 xenografts over time under indicated treatments. Y axis shows the % volume, normalized with initial volume measurements at the start of treatment. n = 4 tumors for PCN and n = 10 tumors for lor and lor+PCN. **I**. Plasma concentrations of brigatinib (brig) from mice treated with indicated drugs for 3.5 weeks. n = 5 mice for all groups. **J**. Change in tumor volume of H3122 xenografts over time under indicated treatments. Y axis shows the % volume as described in panel D. n = 4 tumors for control, n = 6 for JIB-04 and n = 10, 9 for brig and brig+JIB-04. **K**. Plasma concentrations of osimertinib (osi) and sotorasib (soto) from mice treated with indicated drugs (2.5 weeks for osi and 4.4 weeks for soto). n = 5 mice for all groups. **L**. Change in tumor volume of PC9 xenografts for osi and H358 xenografts for soto over time under indicated treatments. Y axis shows the % volume as described in panel H, J. n = 4 tumors for untreated and JIB-04 and n = 10 tumors for osi/soto and osi/soto+JIB-04. For all experiments, dosages used were, 25mg/kg alectinib, 10mg/kg lorlatinib and 55mg/kg JIB-04, 100mg/kg PCN, 20mg/kg brigatinib, 10mg/kg osimertinib and 50mg/kg sotorasib. Mann-Whitney t-test was performed for all concentration data unless specified otherwise and 2-way ANOVA with Sidak’s test for volume data. Mean ± SEM are shown for all, *p < 0.05, **p < 0.01, ***p < 0.001, and ****p < 0.0001.

To discriminate between direct and indirect effects of JIB-04, we examined the impact of JIB04 on CYP3A11 expression *in vitro*, using a mouse hepatocyte cell line, AML12. JIB-04 treatment significantly upregulated CYP3A11 transcript levels, indicating a direct effect **(Fig. S2F)**. Notably, a similar upregulation was observed in a human hepatocyte cell line, Huh7 **(Fig. S2F)**, indicating the broader generalizability of the effect. To assess the functional impact of the elevated CYP3A11 expression, we evaluated the effect of JIB-04 on a well-established blood plasma marker of CYP3A4/CYP3A11 activity, 4β-hydroxy cholesterol to cholesterol (4β-OHC/C) ratio ^26–28^. We found that JIB-04 treatment upregulated 4β-OHC levels and 4β-OHC/C ratio in the blood **(Fig. 3C, S2G)**. To assess whether elevated CYP3A11 activity enhanced alectinib metabolism, we evaluated the impact of JIB-04 on the ratio of alectinib to the product of its conversion by CYP3A4/CYP3A11, the M4 metabolite^29^. As expected, we found that JIB-04 treatment significantly enhanced M4/alectinib ratio, supporting the notion that JIB-04 enhances CYP3A11-mediated metabolic conversion of alectinib **(Fig. 3D)**.

To validate the link between the enhanced CYP3A11 activity and ALKi resistance, we asked whether the resistance-promoting effect of JIB-04 could be phenocopied with a chemically unrelated activator of CYP3A11. Due to the species specificity of CYP regulation, the majority of well-characterized inducers of human CYP3A4 are not potent activators of murine CYP3A11. Therefore, we tested the effect of a known inducer of the murine CYP3A11, pregnenolone 16-alpha carbonitrile (PCN)^25^. As expected, PCN strongly induced CYP3A11 protein expression in mouse liver **(Fig. 3E)**. This enhancement was accompanied by an increase in the plasma marker of CYP3A activity, 4β-OHC/C ratio **(Fig. 3F and Fig. S3A)**. PCN co-treatment dramatically reduced plasma levels of lorlatinib **(Fig. 3G)** and led to a rapid onset of lorlatinib resistance in the H3122 xenograft model **(Fig. 3H)**. These experiments strongly support the notion that pharmacological activation of CYP3A11 can cause resistance to ALKi, alectinib and lorlatinib.

Metabolic elimination of small-molecule drugs typically involves the action of multiple CYP family members. CYP3A4 is not only a primary mediator of metabolic conversion of 30-50% of the current FDA-approved drugs but it is also involved in the metabolic conversion of most drugs^11,18,30^. Therefore, we reasoned that substantial CYP3A11 induction might accelerate PK of drugs that are primarily converted by other CYP family members. To test this assumption, we examined the impact of JIB-04 on plasma levels and therapeutic effects of brigatinib, a clinically approved ALKi inhibitor which is primarily metabolized by CYP2C8, with CYP3A4 being responsible for 27.6% of brigatinib conversion under baseline conditions^31^. Indeed, while sensitivity of H3122 cells to brigatinib was not affected by JIB-04 *in vitro* **(Fig. S3B)**, JIB-04 co-treatment reduced brigatinib blood concentration *in vivo*, translating to reduced responses of H3122 xenograft tumors **(Fig. 3I, J, S3C)**.

Next, we asked whether JIB-04 can cause resistance to other clinically relevant targeted therapy agents that are metabolized by CYP3A4/CYP3A11. To this end, we examined the impact of CYPA311 induction on the sensitivity to other targeted therapies used in NSCLC. Specifically, we assessed the effects of JIB-04 on sensitivity of PC9 xenograft tumor model to EGFR inhibitor, osimertinib, a first-line treatment for EGFR NSCLC, and of H358 xenograft model to sotorasib, the preferred second line therapy in KRAS^G12C^ NSCLC^32^. Notably, JIB-04 did not impact the *in vitro* sensitivity of PC9 and H358 cells to osimertinib and sotorasib, respectively, indicating the lack of a direct, cell-intrinsic effect **(Fig. S3D)**. In contrast, JIB-04 reduced the plasma concentration of both drugs **(Fig. 3K)**. This effect was accompanied by the expected elevation of CYP3A11 protein levels in the liver and a fast onset of resistance of the xenograft tumors **(Fig. 3L, S3E)**. In summary, our results suggest that CYP3A4/CYP3A11 activation might cause resistance to a broad range of therapeutic agents.

### Variability in CYP3A4 activity correlates with variability in clinical outcomes

Therapy resistance is typically attributed to tumor-intrinsic resistance mechanisms. However, recent studies support the potential importance of systemic, PK-related mechanisms: patients treated with the same dose of alectinib can display substantial variability in plasma drug concentrations, with lower drug concentrations correlating with shorter progression-free survival (PFS)^16^. Whereas this variability has not been explicitly linked to altered CYP3A4 expression, multiple pharmacological, dietary, and hereditary factors can alter the expression of CYP3A4 in the liver, which can lead to 10–100-fold variability in the population^12,33^. Therefore, we asked whether variability in CYP3A4 activity could contribute to the variability in clinical responses to therapeutic agents metabolized by this enzyme. To this end, we acquired blood samples from two small cohorts of patients from completed Moffitt-run clinical trials, allowing us to assess the link between the plasma marker of CYP3A4 activity, the 4β-OHC/C ratio and PFS **(Fig. 4A)**. The first cohort contained pre- and post-treatment samples from seven EGFR NSCLC patients^34^ treated with erlotinib, a former first-line therapy for EGFR NSCLC patients and a CYP3A4 substrate^32,35^. The cohort displayed a substantial variability in PFS, ranging from 5.23 to 27.53 months (mean±SD = 13.62±8.85 months) **(Fig. S4A)**. Mass spectrometry-based analyses revealed a substantial inter-patient variability in the 4β-OHC/C ratio (range = 0.11*10^-4^-0.55*10^-4^; mean±SD = 0.29*10^-4^±0.13*10^-4^). Despite a small cohort size, Spearman correlation analyses revealed a significant negative correlation between pre-treatment CYP3A4 activity, and the duration of PFS **(Fig. 4B)**, supporting the notion that higher CYP3A4 activity is expected to contribute to faster onset of resistance. Notably, while the negative correlation trend was also observed in post-treatment blood samples, the effect did not reach statistical significance **(Fig. S4B)**, reflecting changes in CYP3A4 activity over the course of treatment (Fig. S4C).

**Main figure 4:**
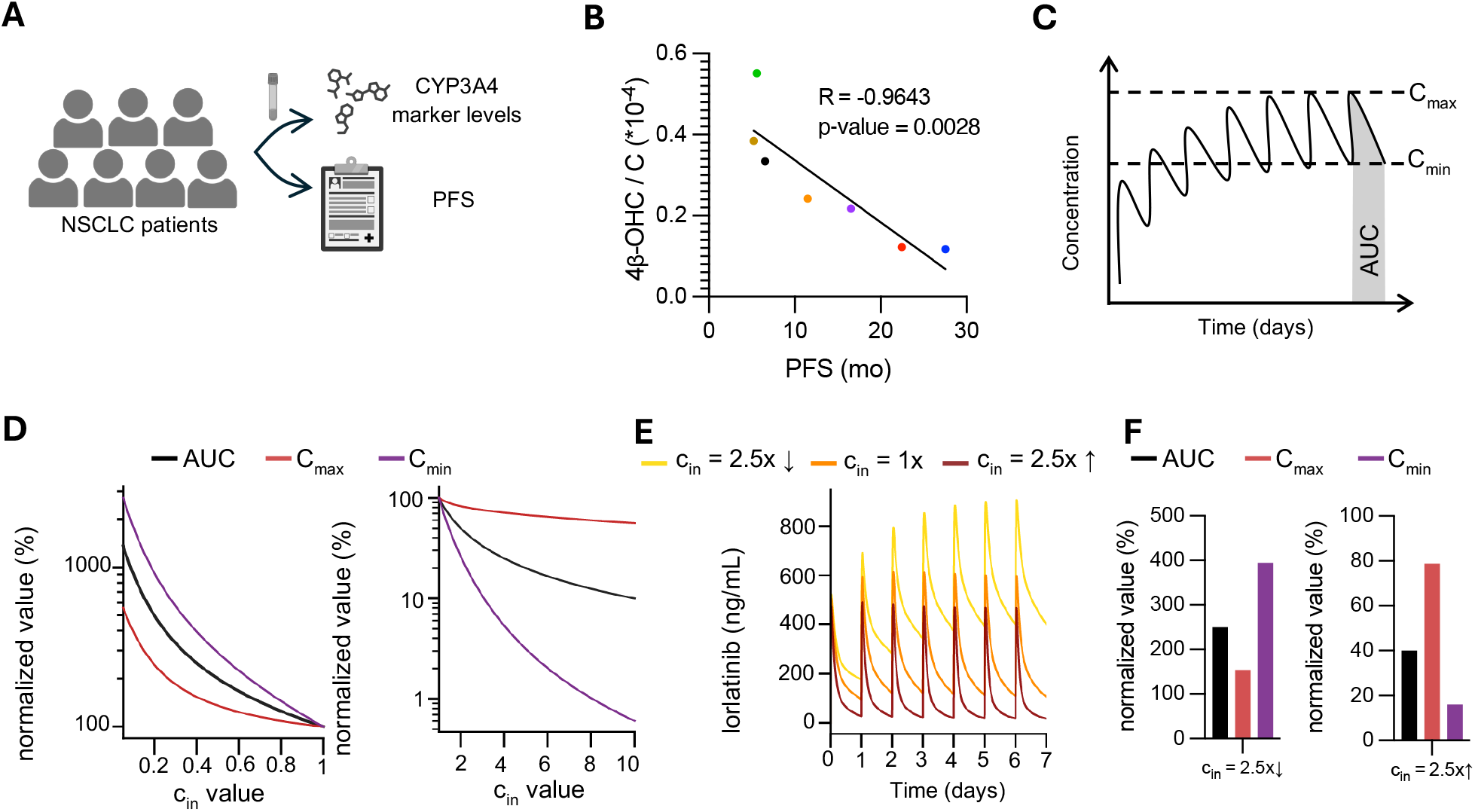
Variability in CYP3A4 activity leads to variability in drug concentrations. **A**. Illustration showing experimental process of patient sample handling. Blood-plasma samples were taken from retrospective studies and CYP3A4 marker was estimated as ratio of quantitation of 4β-OHC/cholesterol. PFS was noted. **B**. Spearman correlation plot with linear regression line is shown between CYP3A4 and PFS. Each color represents an EGFR NSCLC patient (prior to treatment with erlotinib). **C**. Schemata showing PK metrics, maximum concentration (C_max_), minimum concentration (C_min_) and area under curve (AUC) at steady state with multiple dosing of lorlatinib. **D**. Dynamics of PK metrics with lower or higher CYP3A4 activity (c_in_). Values are normalized to that at c_in_ of 1 as percentages. **E**. Concentration-time curves of lorlatinib with standard of care dosage (SOC; 100mg QD) at indicated c_in_ values. **F**. Quantification of PK metrics in E at steady state. Values are normalized to that at c_in_ of 1 as percentages.

The second cohort contained post-treatment samples from thirteen KRAS NSCLC patients treated with sotorasib with variable PFS **(Fig, S4D)**. This cohort included five patients who also had matched blood samples taken at baseline. In contrast to the erlotinib cohort, we did not detect a significant correlation between 4β-OHC/C ratios in pre-treatment samples and PFS **(Fig. S4E, F)**. On the other hand, analyses of blood samples with matched pre-post therapy measurements revealed a substantial elevation in the 4β-OHC/C ratios in four out of five patients **(Fig. S4G)**. Notably, sotorasib is a known CYP3A4 inducer, documented to enhance CYP3A4-mediated drug clearance by ∼50 %^36^. While the sotorasib-induced CYP3A4 activation is assumed to plateau at 22 days of treatment^37^ our result suggests the possibility that continuous induction over the longer time frames might contribute to the eventual sotorasib treatment failure. Further, this activation creep, together with patient-patient variability in the magnitude and timing of induction likely obscures correlation analyses based on pre-treatment samples. Therefore, while we did not observe a significant correlation between baseline CYP3A4 markers and progression-free survival in the sotorasib-treated cohort, these findings remain consistent with our hypothesis when considering the drug’s established clinical pharmacology.

### Impact of CYP3A4 variability on systemic drug levels

To gain a systematic, quantitative understanding of the impact of altered drug metabolism on systemic drug concentrations over time, we employed a mathematical modeling approach. To this end, we focused on the current frontline ALKi lorlatinib, adopting the 2-compartment lorlatinib population pharmacokinetic model fitted using PK data from 425 participants treated with the 100mg QD standard of care (SOC) dosage^37^. We extended the model by introducing a parameter, c_in_, representing a change of CYP3A4 activity relative to an average activity level (c_in_ of 1) (**Fig. S5A, Table S1**). Thus, we considered the impact of variable CYP3A4 activity on three key PK metrics area under curve (AUC) representing total exposure, maximum concentration (C_max_) and minimum concentration (C_min_) once systemic drug concentrations reached steady-state fluctuations, thereby excluding the initial loading phase from our comparative analysis (**Fig. 4C**). Whereas our study so far has focused on the effect of accelerated PK, reduced drug elimination might present an equally serious problem in the clinics due to the enhanced toxicity resulting from the elevated drug levels^38–41^.

We started with the analysis across a broad 100x range of metabolic activity (c_in_ 0.1-10) to understand the consequences of the whole range of the reported variability in CYP3A4 expression^12,33^. Our analyses highlighted asymmetry in responses to both increased and reduced metabolic activity; while altered CYP3A4 activity impacted all metrics, it did so with varying degrees of non-linearity **(Fig. 4D)**. Hypoactivation (c_in_ < 1) led to a global but non-linear elevation of all three metrics, with AUC, C_max,_ and C_min_ rising at different rates. Conversely, hyperactivation led to a global decrease in the three metrics, with similar asymmetry. Notably, the impact of elevated metabolism was more aggressive. For example, while 10x reduction in CYP3A4 activity led to a ∼20-fold increase in C_min_, the 10x increase collapsed C_min_ by over two orders of magnitude **(Fig. 4D)**. This fast decay indicates that C_min_ might be the most labile parameter, creating sub-therapeutic windows that may allow for tumor recovery and facilitate the onset of resistance.

Next, to better ground our analyses in clinical reality, we focused on a more conservative range of 2.5x change in c_in_ in both directions. Notably, this range is within the over 8-fold variability in 4β-OHC/C ratios observed in our sotorasib cohort **(Fig. S4F)**. Incorporation of the temporal dimension of PK highlighted the biological risk of hypermetabolism **(Fig. 4E)**. Beyond the reduction in total exposure, the rapid decay of C_min_ creates distinct sub-therapeutic windows between doses. During these intervals, concentrations fall below the minimum effective threshold, potentially permitting tumor recovery and providing a systemic reservoir for the emergence of resistance. Conversely, in the case of hypometabolism, drug concentrations stay above the C_max_ observed under normal metabolism **(Fig. 4E)**. Consistent with our analyses in **Fig. 4D**, the C_min_ was disproportionally affected by both increase and decrease in metabolic activity **(Fig. 4F)**; however, the magnitude indicates that it can be a major contributor to both systemic resistance and toxicity, respectively. In conclusion, our in-silico analyses indicate that variability in CYP3A4 activity can act as a systemic driver of both toxicity and resistance, governed by a non-linear relationship that destabilizes the therapeutic window under standard dosing regimens.

### Therapy resistance from accelerated PK can be mitigated through pharmacological and dose adjustment strategies

In an ideal-case scenario, normal therapeutic window might be restored by identification and exclusion of CYP3A4 modulators from dietary inputs and orthogonal medications^11,43^. However, due to the complex nature of CYP3A4 regulation, including a substantial heritable component, this solution might not be easily available. Therefore, we explored alternative strategies for the mitigation of the effects of altered PK.

Taking advantage of capturing the impact of altered CYP3A4 in the formalism of mathematical models, we started by exploring mitigation strategies, involving adjusting dosage and frequency of drug administration. As a case in point, we focused on 2.5 fold up or downregulation of CYP3A4 activity, starting with exploring the impact of adjusting dosing. While adjusting the dose to match the baseline values for each of the three parameters can be readily achieved under both increased and decreased CYP3A4 activity (horizontal dashed lines in **Fig. 5A**), the required dose differs for each of the three parameters. For example, dose adjustment, required to match the baseline overall drug exposure (vertical dashed lines in **Fig. 5A**) leads to the C_max_ and C_min_ values that are substantially different from the baseline values (**Fig. 5A**). For the reduced metabolism, dose adjustment to 40 mg QD, required to match baseline AUC reduces of the difference between C_max_ and C_min_, with both parameters staying within the normal therapeutic range (**Fig. 5A, B**). In contrast, dose, dose increase to 250 QD, required to return the AUC to the baseline levels under 2.5 fold increased CYP43A4 activity, increases the difference between C_max_ and C_min_. Under 250 mg QD, C_max_ is increased to ∼2x higher value than baseline, while C_min_ was still ∼2.5 fold below the baseline value (**Fig. 5A, C, Table S2**). Therefore, while dose adjustment strategy offers a straightforward solution for mitigating reduced drug metabolism, the utility of this approach is more limited for accelerated metabolism.

**Main figure 5:**
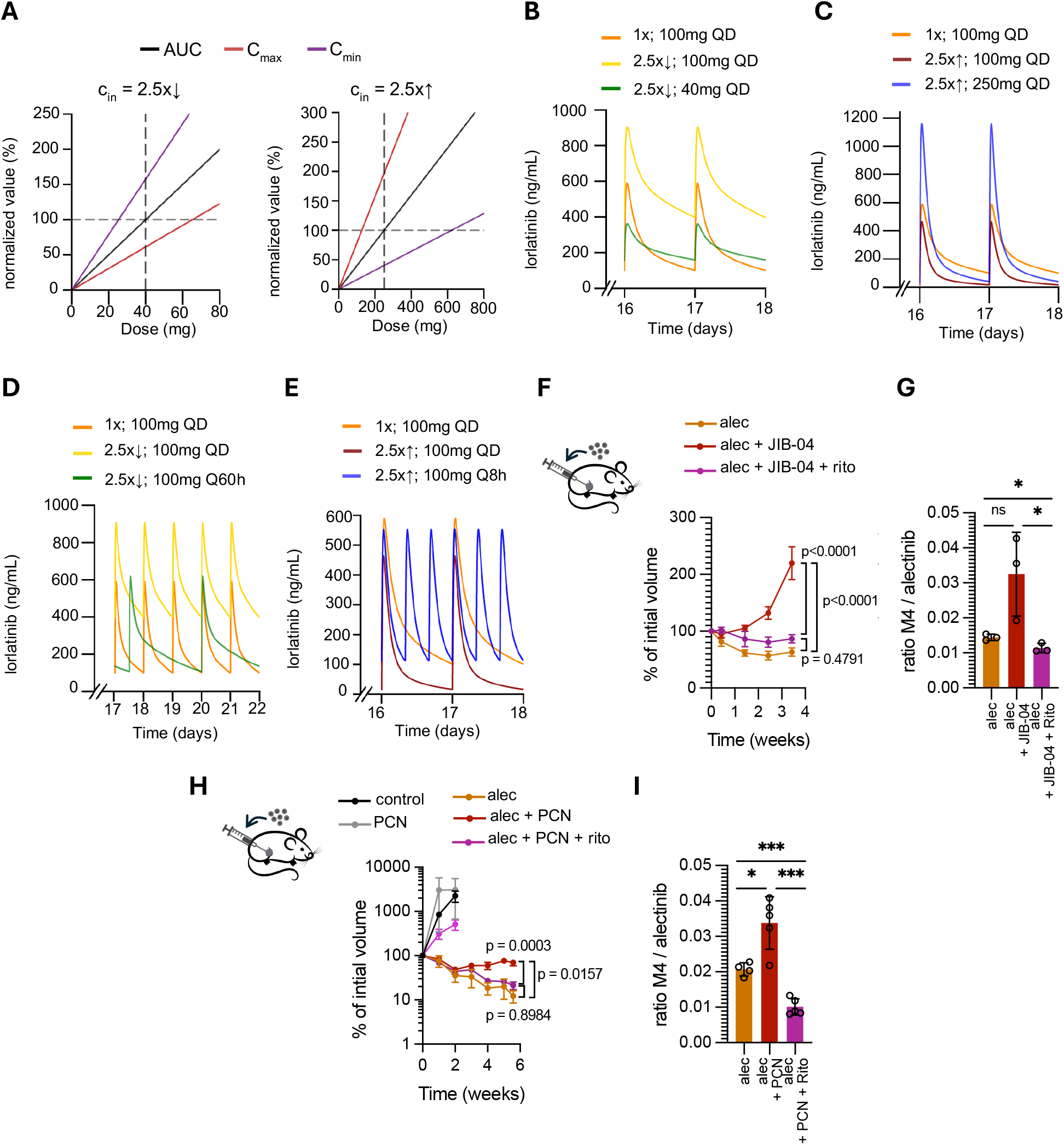
Mitigation strategies for altered CYP3A4/11activity. **A**. Dynamics of PK metrics with change in dosages at indicated CYP3A4 activity (c_in_). Values are normalized as percentage to that at c_in_ of 1 with SOC dosage/interval. **B-E**. Concentration-time curves of lorlatinib at indicated c_in_ values and dosage/interval (noted as c_in_ ; dosing). **F**. Change in tumor volume of H3122 xenografts over time under indicated treatments. Y axis shows the % volume, normalized with initial volume measurements at the start of treatment. n = 5 for alec and 6 tumors for rest. 25mg/kg alectinib, 55mg/kg JIB-04 and 50mg/kg ritonavir (rito). 2-way ANOVA with Sidak’s test was performed. **G**. M4/alec ratio in blood collected from mice treated with indicated drugs for 3.5 weeks. n = 3 mice for all groups. Unpaired t-test was performed. **H**. Change in tumor volume of H3122 xenografts over time under indicated treatments. Y axis shows the % volume, normalized with initial volume measurements at the start of treatment. n = 9, 10 tumors for alec + PCN and alec + PCN + rito, n = 8 for alec and n = 6 for control, PCN and rito. **I**. M4/alec ratio in blood collected from mice treated with indicated drugs for 5.5 weeks. n = 4 mice for alec and n = 5 mice for rest. Unpaired t-test was performed. 25mg/kg alectinib, 100mg/kg PCN and 50mg/kg ritonavir. 2-way ANOVA with Sidak’s test was performed. Mean ± SEM are shown for all, *p < 0.05, **p < 0.01, ***p < 0.001, and ****p < 0.0001.

Next, we used our PK model to explore the utility of adjusting frequency of drug administration. Whereas the dosing frequency adjustment required to achieve a match with the baseline levels differed for the three key PK parameters, the differences were less pronounced compared to the dose adjustment strategies (**Fig. S5B**). For the 2.5x reduction in CYP3A4 activity, adjusting the frequency of drug administration from 24 hours to 60 hours (Q60h) produced a close match to the baseline (**Fig 5D**). Whereas this administration frequency is not practical, a similarly close match could be achieved with doubling the drug administration frequency to 48 hours, while reducing the dosage to 76 mg (**Fig. S5C**). Dose adjustment also provided an adequate solution for the 2.5x increase in CYP3A4 activity, as shift to drug administration to every 8 hours brought all three parameters within a 10% window of the baseline values (**Fig. 5E**). In summary, apart from practical compliance considerations of dose interval adjustments, it provides a superior strategy for mitigating the consequences of increased and decreased drug metabolism within the 2.5x change range.

Next, we explored the suitability of dose and frequency mitigation strategies for a higher change in the CYP3A4 activity levels, ∼10-fold increase in CYP3A4 activity, consistent with our experimental data in mice and that reported with strong CYP3A4 inducers^44^. Thus, we tested the impact of c_in_ of 10 on concentrations and found that while C_max_ only reduces by 44%, the C_min_ reduces drastically by 99% from 100.8 ng/mL to 0.6 ng/mL **(Table S2)**. Dosing adjustment of 1000mg QD to rescue PK metrics can have substantial toxicity concerns (data not shown) and interval adjustments of 100mg Q2.4h for oral dosing in unfeasible **(Fig. S5D)**. Additionally, the relatively feasible combined dosing and interval solution of 200mg Q6h rescues AUC but exhibits high fluctuations in concentrations, without rescuing C_max_ and C_min_ within 10% of SOC levels **(Table S2)**. We speculate that using alternative routes of drug administration such as continuous infusion should be able to provide optimal and stable systemic concentrations of the drug in cases of strong CYP3A4 hyperactivation.

Finally, we explored an alternative solution to strong CYP3A4 activation. We asked whether co-treatment with a CYP3A inhibitor may provide clinically feasible and readily applicable solution to CYP3A4 mediated resistance for strong CYP3A4 activation. We tested ritonavir, a widely used pharmacological enhancer that boosts the effect of antiretroviral medications in several clinical contexts by inhibiting CYP3A enzymes, including human CYP3A4 and murine CYP3A11^45^. Ritonavir co-treatment counteracted the effects of JIB-04, enhancing the sensitivity of tumors to alectinib **(Fig. 5F)**. Further, ritonavir reduced 4β-OHC/C and M4/alec ratio in the blood in JIB-04-treated mice, indicating the expected inhibition of the CYP3A11 activity **(Fig. 5G, S5F, G)**. We observed similar rescuing of sensitivity with PCN as well **(Fig. 5H, I, S5H, I)**.

Overall, our experimental and mathematical model results indicate that dose and interval adjustments can help rescue concentrations under baseline CYP3A4 variability, however, strong CYP3A4 hyperactivation may require pharmacological intervention to alleviate the impact of accelerated drug metabolism.

## Discussion

Our study suggests that accelerated drug metabolism, mediated by increased expression of CYP3A4 and related P450 enzymes, can lead to resistance to targeted therapies. Despite a general recognition of the importance of PK considerations and the detailed understanding of the molecular mechanisms that underlie drug metabolism, accelerated PK is not commonly considered to be a potential resistance mechanism in clinical practice. Importantly, this resistance mechanism might be relatively easy to address by monitoring/adjusting dietary intake and other therapeutics, using pharmacological modulators of CYP4A4 activity or by modeling-guided adjustment of dosing and frequency of therapy administration. While our study has focused on the context of cancer-targeting therapies, we expect PK-mediated resistance and its mitigation strategies to be relevant to any type of small-molecule therapies, metabolized by CYP3A4 and P450 enzymes in general.

Both intrinsic and acquired therapy resistance are typically attributed to tumor-intrinsic mechanisms that reduce the sensitivity of tumor cells to drug exposure. These include mutational mechanisms that amplify the target or impair drug binding, mutational and epigenetic mechanisms that activate bypass signaling, enhanced drug efflux, etc.^2–4,13,46^. Further, there is a growing recognition of the potential importance of microenvironmental interactions, as multiple paracrine and juxtacrine mechanisms have been shown to reduce tumor cell sensitivity by activating alternative signaling pathways and inducing phenotypic plasticity and the involvement of its various components in resistance. These include paracrine interactions with immune cells, fibroblasts, extracellular matrix, etc., through cytokines, growth factors, and metabolites^3,5,6,10,13,47^. Still, in a significant fraction of clinical cases, a known resistance mechanism cannot be identified. Further, most studies and clinical case reports attribute resistance to a single cell intrinsic or microenvironmental mechanism. However, recent findings from our lab and other groups suggest a more complex scenario of a multifactorial therapy resistance, which integrates multiple cell intrinsic (including mutational and epigenetic changes) and microenvironmental mechanisms^10,13,48^. Therefore, even in those cases where a known resistance mechanism is implied, accelerated PK might still represent a substantial contributor to therapy failures.

Throughout different targeted therapy contexts, populations of patients display substantial variability in clinical responses. This variability likely reflects inter-tumor heterogeneity, as each individual cancer represents a result of a unique evolutionary trajectory^1,49^. At the same time, the impact of inter-tumor heterogeneity is not mutually exclusive with the variability in drug bioavailability. Orally administered drugs are subject to the absorption, distribution, metabolism, and excretion dynamics. Therefore, variability in expression of drug transporters in the gastrointestinal system, gastric pH, specific food intake, and kidney function, etc., should be expected to contribute to the variability in clinical outcomes^21^. Indeed, recent studies of cohorts of ALK+ NSCLC documented substantial inter-patient variability in plasma alectinib levels, in turn reflected in the variability in clinical outcomes^16^. Variability in drug metabolism due to inter-patient differences in the expression of metabolic enzymes might be particularly important. Similar to xenobiotics that are introduced into the circulation with food intake, small-molecule drugs are metabolized by P450 family enzymes, altering their activity and converting them into more soluble metabolites, thereby facilitating excretion^18,22^.

CYP3A is one of the key P450 family members, involved in the metabolism of majority of oral small molecule drugs, including FDA-approved anti-cancer targeted therapies^11,18,30^. Importantly, expression and activity of P450 enzymes, including CYP3A4 can be augmented and suppressed by a broad spectrum of xenobiotics, including compounds in herbal supplements (Yin Zi Huang, Echinacea purpurea, Ginkgo biloba, St. John’s wort etc.) and many drugs (such as benzodiazepines, HIV antivirals, calcium channel blockers, statins, antibiotics etc.)^22,43^. These compounds can inhibit P450 enzyme activity via direct binding or activate it by inducing gene expression or protein stabilization^43^. Additionally, expression of CYP3A4 is influenced by multiple genetic polymorphisms that increase or reduce its expression^30^. Consequently, studies documented variability in CYP3A4 activity that spans several orders of magnitude^12,33^.

Detailed PK evaluation constitutes a mandatory element of the clinical drug development pipelines^50^. However, this evaluation is limited to preclinical phase and stage 1 clinical trials, which rely on carefully selected, small groups of patients that might not adequately recapitulate PK variability in larger patient cohorts. Consistent with the central role of CYP3A4 in drug metabolism, our preclinical studies reveal that enhanced CYP3A4 activity leads to strong resistance to multiple targeted therapy inhibitors. Notably, enhanced CYP3A4 activity could be expected to enhance the metabolic breakdown of drugs that are primarily metabolized by other P450 family enzymes. Even though at normal conditions only 27.6% of brigatinib is known to be metabolized by CYP3A4^31^, induced CYP3A activity caused brigatinib resistance in xenograft models of ALK+ NSCLC **(Fig. 3J)**. Importantly, CYP3A4 expression and activity can vary over time due to dietary changes and treatment with other drugs. Moreover, our analyses of the sotorasib-treated cohort indicate a possibility of autoinduction of metabolism for some of the drugs over longer time frame, likely contributing to the therapy resistance **(Fig. S4G)**. Whereas our study has focused on enhanced CYP3A4 activity, reduced activity might also represent a substantial clinical challenge, as elevated drug levels resulting from inhibition of drug metabolism can be expected to increase systemic toxicities, which might force termination of potentially life-saving treatments^39–42^.

Assessing the true extent of the clinical relevance of accelerated PK, mediated by enhanced CYP3A4 activity, would require robust analyses of larger cohorts. Still, both first principles considerations and our pilot analyses indicate that it is likely to represent a substantial contributor to clinical therapy resistance. Targeted therapies are given as a one-size-fits-all standard dose for all patients irrespective of their metabolic function. However, studies have reported 24-84% inter-patient variability in PK exposure to oral targeted therapies where some patients may develop adverse events due to high drug exposure, while others may experience lack of therapeutic efficacy. About 30% patients have been reported to be underdosed, and about 15% overdosed^41,42^. In contrast to multifaceted tumor cell intrinsic resistance mechanisms, therapy resistance stemming from accelerated drug metabolism should be relatively easy to detect. Direct measurements of plasma drug concentrations are complicated by the variability in food intake, temporal dynamics of drug PK and the need for the use of drug-specific tests. In contrast, CYP3A activity might be easier to assess in blood plasma samples. Our study used 4βOHC/C, a well-established marker for the activity of CYP3A4 and CYP3A5^26–28^. While the long half-life of 4βOHC could be limiting in the contexts of shorter time scale therapeutic contexts, this is less of a problem for cancer-targeted therapies that are often administered over the course of multiple months and even years. While 4bOHC/C might not necessarily provide a robust proof of the elevated CYP3A4 activity, its non-invasive nature and reasonable accuracy make it a great candidate for an initial screen, which can be followed up with a more direct and quantitative interrogation of CYP3A4 activity using more direct methods, such as using CYP3A4 substrate midazolam^28^.

Importantly, therapy resistance and increased systemic toxicity resulting from enhanced and reduced CYP3A activity could be more amenable to mitigation compared to addressing divergent tumor-intrinsic mechanisms. Given that P450 activity can be influenced by multiple dietary components, xenobiotics from food supplements and pharmacological drugs, the obvious first solution could be a careful examination of the patient’s diet and co-medications to identify and exclude P450 modulators. The model-based personalized adjustment of both dosing and frequency of therapy administration can provide a good generalizable solution for baseline CYP3A4 activity, returning systemic drug concentrations to the desired range. In those cases where this intervention is unsuccessful, the effects of accelerated CYP3A4 activity could be potentially mitigated by pharmacological interventions, such as the use of a pharmacological inhibitor, ritonavir.

## Methods

### Cell lines

The cells lines, H3122, STE-1, PC9 and H358 were received from the Lung Cancer Center of Excellence cell line depository at the Moffitt Cancer Center. These cell lines were cultured in RPMI (Gibco) supplemented with 10% fetal bovine serum (Gibco), 1% penicillin/streptomycin (Gibco) and 10μg/mL human recombinant insulin (Gibco). AML12 cells were purchased from ATCC and cultured as per recommended handling protocol (DMEM F12 (Gibco), 10μg/mL insulin, ITS (Corning) and 40ng/mL dexamethasone (AdipoGen) along with 1% penicillin/streptomycin). Huh7 cells were received as a gift from Dr. Ken Wright at Moffitt Cancer Center and maintained in DMEM F12 media supplemented with 10% FBS and 1% penicillin/streptomycin. All cell lines were maintained at 37°C and 5% CO_2_.

mCherry variants of H3122 cells were generated as described previously^51^. For Gluc expressing H3122 cells, Gaussia luciferase (GLuc) plasmid (pMCS-Gaussia Luc, Thermo Fisher Scientific) was cloned into pLenti6.3/V5 as described previously^47^.

### In vitro assays

For the cell titer glo of evolving and evolved resistant cells, the H3122 cells were grown in 2μM alectinib, 2μM lorlatinib or 0.5μM crizotinib for 3 weeks (evolving cells) and evolved resistant cells were derived as previously described^13^. 2000 cells / well were seeded in 96 well white clear bottom plates and treated with 1uM JIB-04 next day. Cell titer glo assay was developed according to manufacturer’s protocol 5 days after treatment. Naïve cells (without ALKi pre-treatment) treated with DMSO was used as control to calculate % viability. For all other cell titer glo experiments, 4000 cells / well were seeded and cell titer glo assay was performed after 72 hours of treatment. DMSO vehicle control was used to calculate % viability. For 3D assays, H3122 or STE-1 cells were seeded at a density of 10,000 cells / well in 3D spheroid ultra-low attachment round-bottom well plates (Corning #4520). Cells were cultured under standard conditions until compact spheroids were formed (∼10 days). IncuCyte (image as well as fluorescence) signal was recorded before treating the spheroids with the mentioned treatment (alectinib, lorlatinib, and/or JIB04). The spheroids were replenished with fresh media every week before adding the treatment to each well. The fluorescence signal was recorded twice a week. The data represents triplicates of each condition, where each time point is normalized to the baseline signal.

### In vivo models

0.5-1 million ALK+ H3122, ALK+ STE-1, EGFR mutant PC9 or KRAS mutant H358 cells were injected subcutaneously in two abdominal flanks of NSG mice. Tumors were measured using vernier caliper once a week as described previously^13^. For the lung tumor model, 100,000 H3122 cells expressing GLuc were injected in the tail vein of NSG mice. To monitor tumor growth, secreted luciferase assay was performed as described separately (supplementary methods).

0.5% HPMC with 0.1% Tween80 was used as vehicle for alectinib, lorlatinib, sotorasib and osimertinib. 12.5% cremaphore (EMD Millipore) with 12.5% DMSO was used as vehicle for JIB-04 and sesame oil for PCN. All drugs used were given as oral gavages daily, except JIB-04 was given twice a week.

### PK analysis

NSG mice were given oral gavage with 55mg/kg of JIB-04 twice a week for 3 doses. JIB-04 pre-treated and untreated mice were starved overnight and subjected to 10mg/kg lorlatinib treatment, given as a single oral dose. Blood samples were collected at multiple timepoints after lorlatinib treatment, 0.25, 0.5, 0.75, 1, 1.25, 2, 4 and 6 hours. Lorlatinib concentrations were measured in plasma.

### Drug and metabolite quantification

Drugs were measured in either tumors or blood-plasma and metabolites were measured in blood-plasma. To prepare tumors for quantification, tumors were snap-frozen at the time of collection and 0.1mg was used. For blood-plasma, blood was collected from either tail vein incision during experiment or using cardiac puncture at the endpoint into heparin coated tubes. Blood was spun at 2000 g for 10 minutes and plasma was collected and stored at -80 until used. Drugs, alectinib, M4 (metabolite of alectinib), lorlatinib, osimertinib, sotorasib and brigatinib and, endogenous metabolites, 4β-hydroxy cholesterol and cholesterol were measured. Plasma was diluted ∼4-fold in PBS for quantification, and the final concentrations were adjusted with the dilution factor. Drugs were quantified at the Pharmacokinetics and Pharmacodynamics core facility and endogenous metabolites were quantified at the Metabolomics core facility, both at Moffitt Cancer Center. Detailed methods are in supplementary methods.

### Western blot

Protein levels were assessed in mouse liver samples. Briefly, livers were snap-frozen at the time of collection and stored at -80°C. For protein extraction, frozen livers were crushed into powder using mortar and pestle while placing on dry ice and pouring liquid nitrogen to keep the tissue cold. Powder was dissolved in RIPA lysis buffer (Fisher, Cat# BP115-500), supplemented with protease and phosphatase inhibitor cocktail (Thermo Scientific, Cat# 78440). Protein was harvested, followed by measurement using BCA assay (Fisher, Cat# 23227) as per manufacturer’s recommendations. Protein expression was assessed using NuPAGE gels (Thermo Fisher) as per manufacturer’s protocol. CYP3A4 antibody (Cell Signaling Technology, Cat# 13384, Clone# D9U6N) and b-Actin antibody (Santa Cruz, Cat# sc-47778) were used at 1:1,000 dilution. HRP conjugated secondary antibodies, anti-mouse (Cell Signaling Technology, Cat# 7076P2) and anti-rabbit (Cell Signaling Technology, Cat# 7074P2) were used at 1:10,000 dilution. HRP chemiluminescent substrate was purchased from Millipore and images were taken using Amersham Imager 600 (GE Healthcare Life Sciences).

### Patient information and analysis

Plasma samples collected in two patient cohorts were used retrospectively and subjected to quantification of 4β-hydroxy cholesterol and cholesterol as described above. The time to disease progression was noted. First cohort included 7 patients with EGFR NSCLC treated with erlotinib (NCT01859026)^34^. Pre- and post-treatment plasma samples were used for all patients. Second cohort included 13 KRAS NSCLC patients treated with sotorasib. Pre-sotorasib treatment samples were available and used for 5 patients with matched post-sotorasib treatment and post-sotorasib treatment samples were used for all other patients.

### Statistics

Unpaired 2-tailed t test with or without Welch correction and Mann-Whitney tests were performed wherever indicated. For tumor volume data comparing multiple groups, 2-way ANOVA was used with multiple comparisons and Sidak’s test. Spearman correlation with linear regression was performed for patient data. All data is represented as mean ± SEM. p < 0.05 was considered significant. Statistical analyses were performed with GraphPad Prism v9.0 (GraphPad Software).

### Study approval

All animal experiments followed IACUC approved protocols. Animals were housed in the vivarium at Moffitt Cancer Center in controlled environment on a 12-hour light/dark cycle with food and water, except in pharmacokinetic experiments where the mice were starved overnight. Patient samples used were banked at the Histology Core facility at Moffitt Cancer Center from consented retrospective studies. Samples were obtained after IRB approval and without patient identification information.

## Data availability

The raw values of the data presented in the study is provided in the Supporting Data Values file. Mathematical model code is available on GitHub at https://github.com/pelesko21/PK_PD_Lorlatinib.

## Acknowledgements

This work has been supported in part by the Proteomics & Metabolomics, Cancer PK/PD, Tissue, Vivarium and the Analytic Microscopy Core Facilities at the H. Lee Moffitt Cancer Center & Research Institute, an NCI designated Comprehensive Cancer Center (P30-CA076292).

