## Supplementary Figures for "Pharmacokinetic Acceleration via CYP3A4 Hyperactivation as a Clinically Actionable Mechanism of Targeted Therapy Resistance in NSCLC"

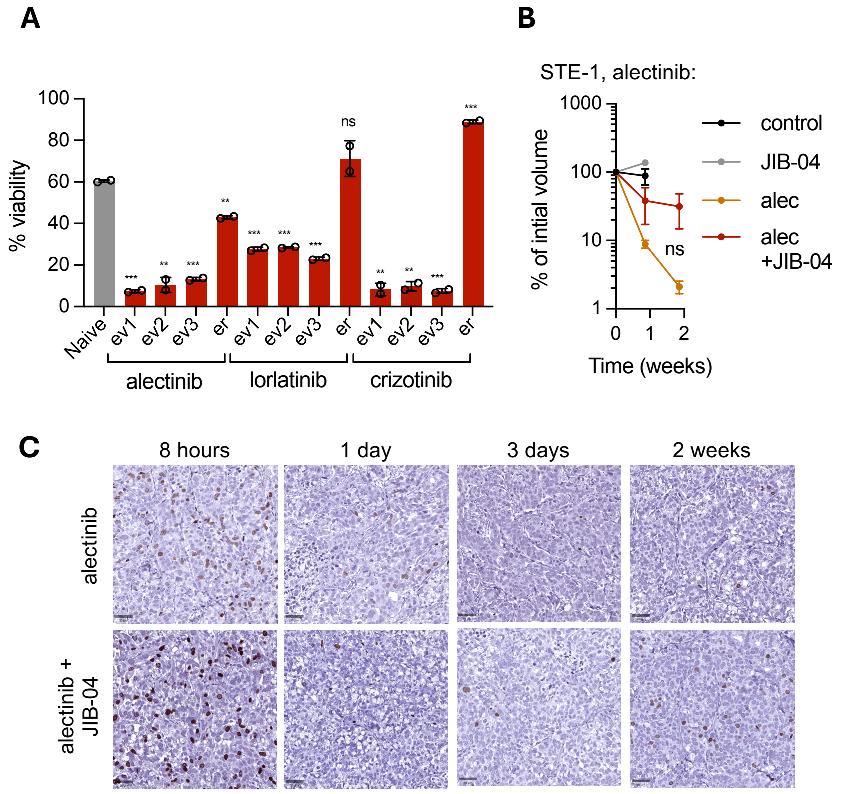


**Supplementary Figure 1: Continued treatment with JIB-04 sensitizes cells to ALKi *in vitro,* while causing resistance *in vivo***

**A.** Cytotoxicity assay of H3122 cells either naïve, pre-treated with 2μM of alectinib (alec) or 2μM of lorlatinib (lor) or 0.5μM of crizotinib (criz) for 3 weeks (ev) or cells with evolved resistance (er) to these drugs. Cells were treated with 1μM of JIB-04 for 6 days. Data shows % viability relative to DMSO control. n = 2 replicates. 3 independent pre-treated lines are shown for 3-week pre-treatment cells. One-way ANOVA with Brown-Forsythe and Welch test was performed. Mean ± SEM are shown. **B.** Change in tumor volume of STE-1 subcutaneous xenografts over time under indicated treatments (25mg/kg alec, 55mg/kg JIB-04). n = 2 tumors for control, n = 4 tumors for JIB-04, alec and alec+JIB-04. p-value represents interaction of 2-way ANOVA analysis with Sidak's test. Mean ± SEM are shown. **C.** Histology images of H3122 xenografts treated with indicated treatments and stained with BrDU (scalebar at 50μm, scanned at 20X) are shown. Tumors were collected at indicated duration of treatment. *p < 0.05, **p < 0.01, ***p < 0.001, and ****p < 0.0001

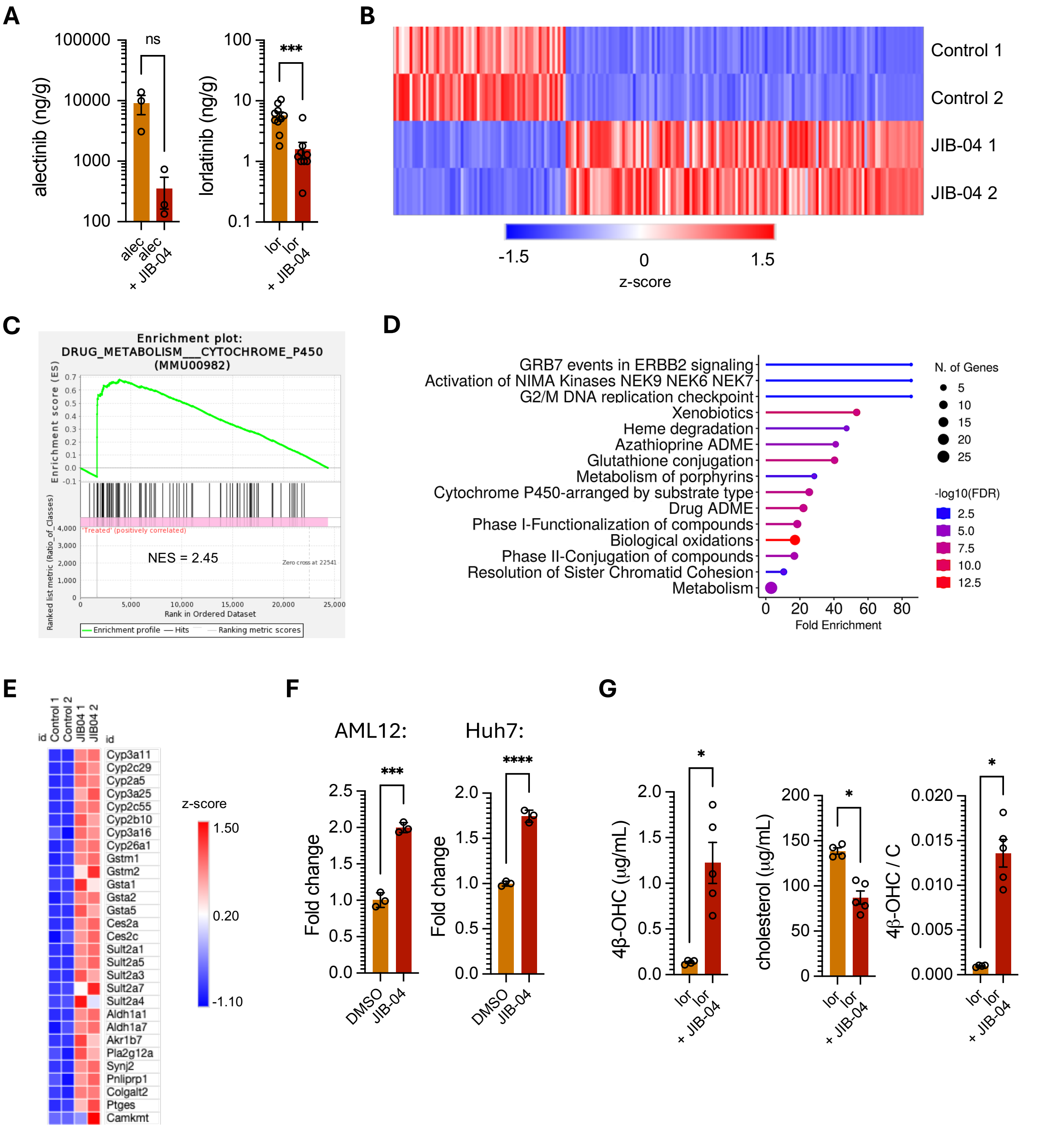


**Supplementary Figure 2: JIB04 increases drug metabolism via CYP3A11 in livers**

**A.** Concentrations of alectinib and lorlatinib in tumors from mice bearing subcutaneous xenograft tumors and treated with indicated drugs for 2.5 weeks for alectinib (n = 3 mice for both groups) and 3.7 weeks for lorlatinib (n = 10 mice for both groups). Mice were sacked 6 hours after last dose and drug concentrations were quantified using LC-MS/MS. Mann-Whitney test was performed. **B.** Heatmap showing z-scores of FPKM of genes with 1.5<log2 fold change<-1.5 derived from differential gene expression analysis (p-value<0.05). **C.** Representative enrichment plot of GSEA pathway with p-adjusted <0.05. **D.** Top 15 Reactome pathways using ShinyGO. Genes used have 1.5<log2 fold change in JIB-04 vs control. **E.** Heatmap showing z-scores of FPKM of genes that were represented in metabolism-related pathways in Fig. 3A. **F.** CYP3A11 and CYP3A4 expression in murine AML12 and human Huh7 hepatocytes treated with 2.5μM and 1μM JIB-04 respectively. Expression was measured by qPCR, and JIB-04 values were

normalized to DMSO controls to obtain fold change. n = 3 replicates. Unpaired t-test was performed. **G.** Plasma concentrations of 4β-OHC, cholesterol and ratio of 4β-OHC with cholesterol. Blood was collected from mice treated with indicated drugs (10mg/kg lorlatinib and 55mg/kg JIB-04) for 5.5 weeks. n = 4 mice for lor and n = 5 mice for lor+JIB-04. Mann-Whitney test was performed. Mean ± SEM are shown for all graphs, *p < 0.05, **p < 0.01, ***p < 0.001, and ****p < 0.0001.

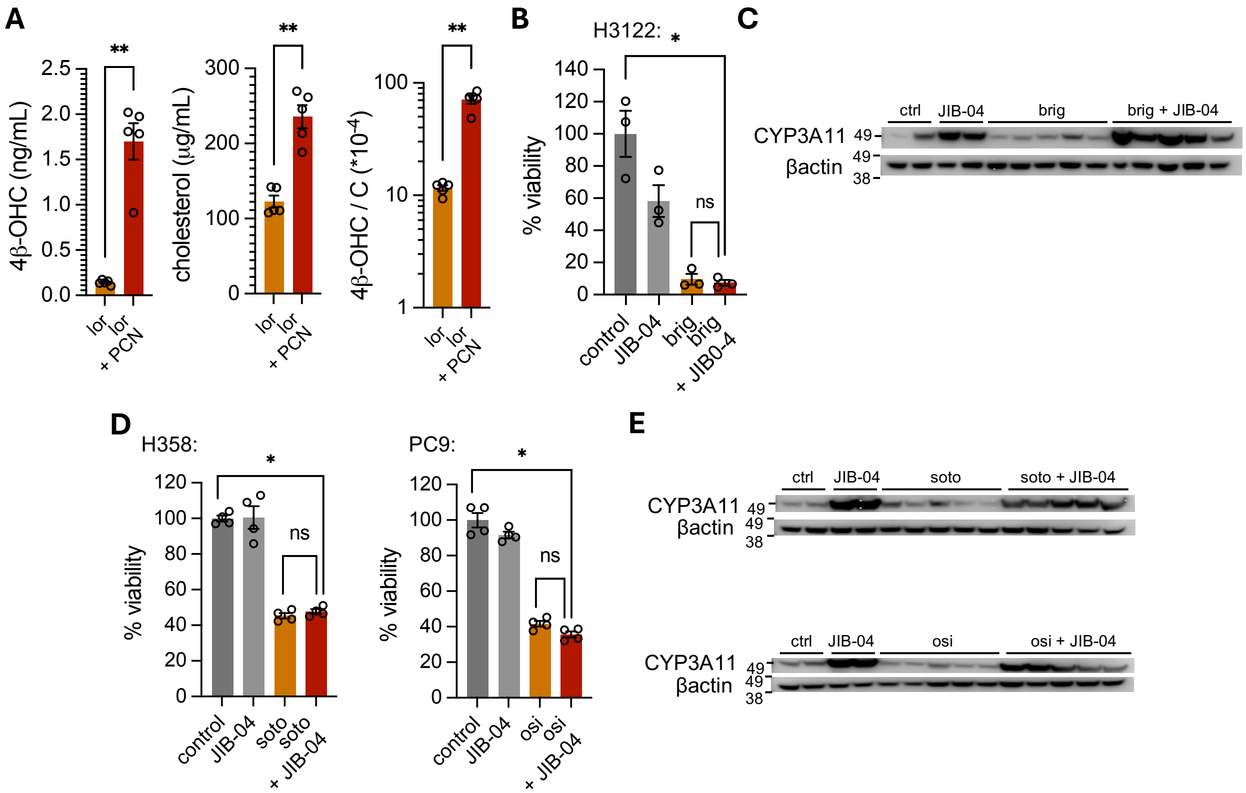


**Supplementary Figure 3: Impact of JIB-04 on multiple targeted therapies**

**A.** Plasma concentrations of 4β-OHC and cholesterol and ratio (4β-OHC/C). Blood was collected from mice treated with indicated drugs for 3.5 weeks. n = 5 mice for all groups. Mann-Whitney test was performed. **B.** Short-term cell titer glow assay for sensitivity of H3122 cells to 0.5µM brigatinib and 2.5µM JIB-04 with DMSO as control. Quantification was performed after 72 hours of treatment. Welch test was performed. **C.** Western blot images showing protein levels of CYP3A11 with βActin as loading control. Proteins were harvested from mice treated with indicated drugs (20mg/kg brigatinib and 55mg/kg JIB-04) for 3.5 weeks. **D.** Short-term cell titer glow assay for sensitivity of H358 cells to sotorasib (0.05µM) and PC9 cells to osimertinib (0.05µM). DMSO was used as control, and inhibitors were combined with 2.5µM JIB-04 as indicated. Quantification was performed after 72 hours of treatment. Mann-Whitney test was performed. **E.** Western blot images showing protein levels of CYP3A11 with βActin as loading control. Proteins were harvested from mice treated with 50mg/kg sotorasib (4.4 weeks) or 10mg/kg osimertinib (2.5 weeks) with 55mg/kg JIB-04 as indicated. Mean ± SEM are shown for all, *p < 0.05, **p < 0.01, ***p < 0.001, and ****p < 0.0001.

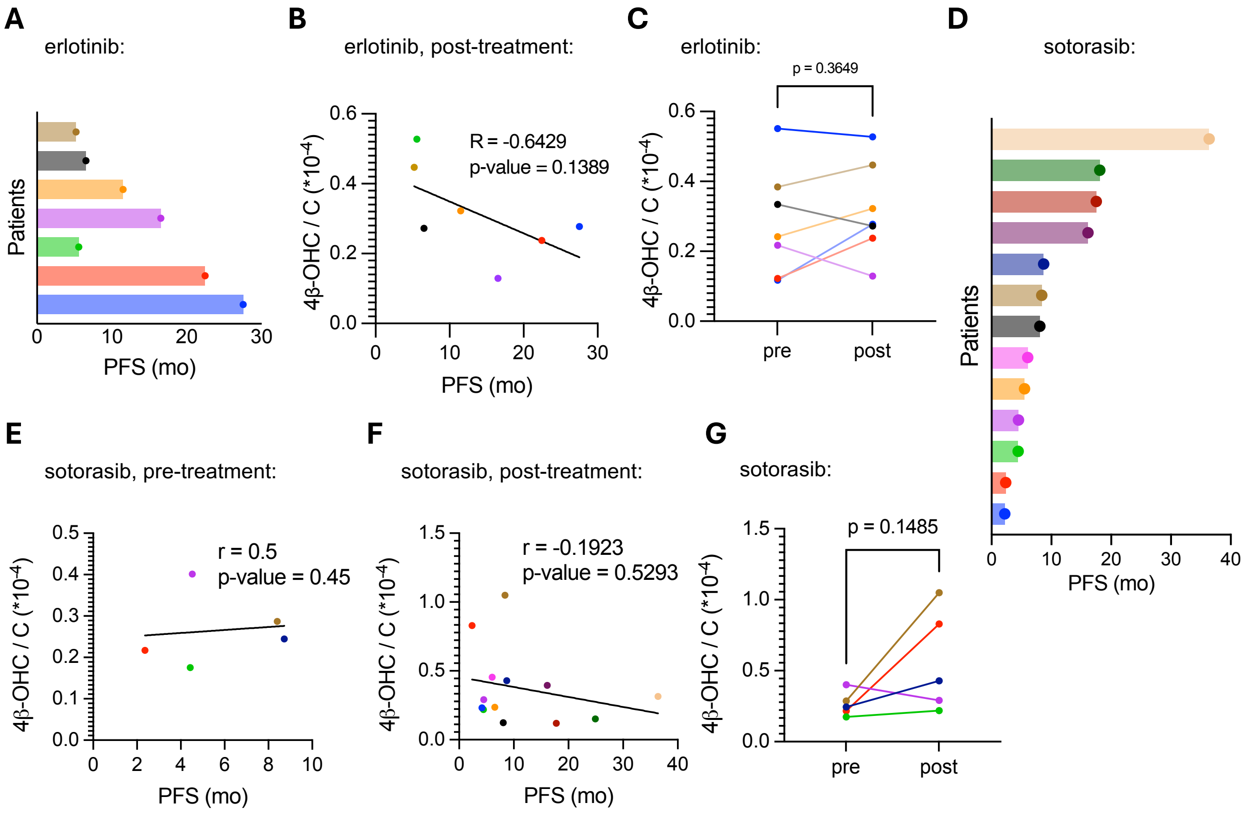


**Supplementary Figure 4: PFS and CYP3A4 activity in erlotinib and sotorasib treated patients**
**A.** Distribution of PFS for EGFR NSCLC patients. **B.** Spearman correlation plot with linear regression line is shown between CYP3A4 activity in post- treatment samples and PFS. Each color represents an EGFR NSCLC patient. **C.** Matched CYP3A4 activity for patients pre- and post- erlotinib treatment. Paired t-test was performed, n = 7 patients. **D.** Distribution of PFS is shown for all sotorasib treated KRAS patients. **E, F.** Spearman correlation plot with linear regression line is shown between CYP3A4 activity and PFS in pre-treatment samples in **E** and post-treatment samples in **F**. Each color represents a KRAS NSCLC patient. **G.** Matched CYP3A4 activity for patients pre- and post- sotorasib treatment. Paired t-test was performed, n = 5 patients.

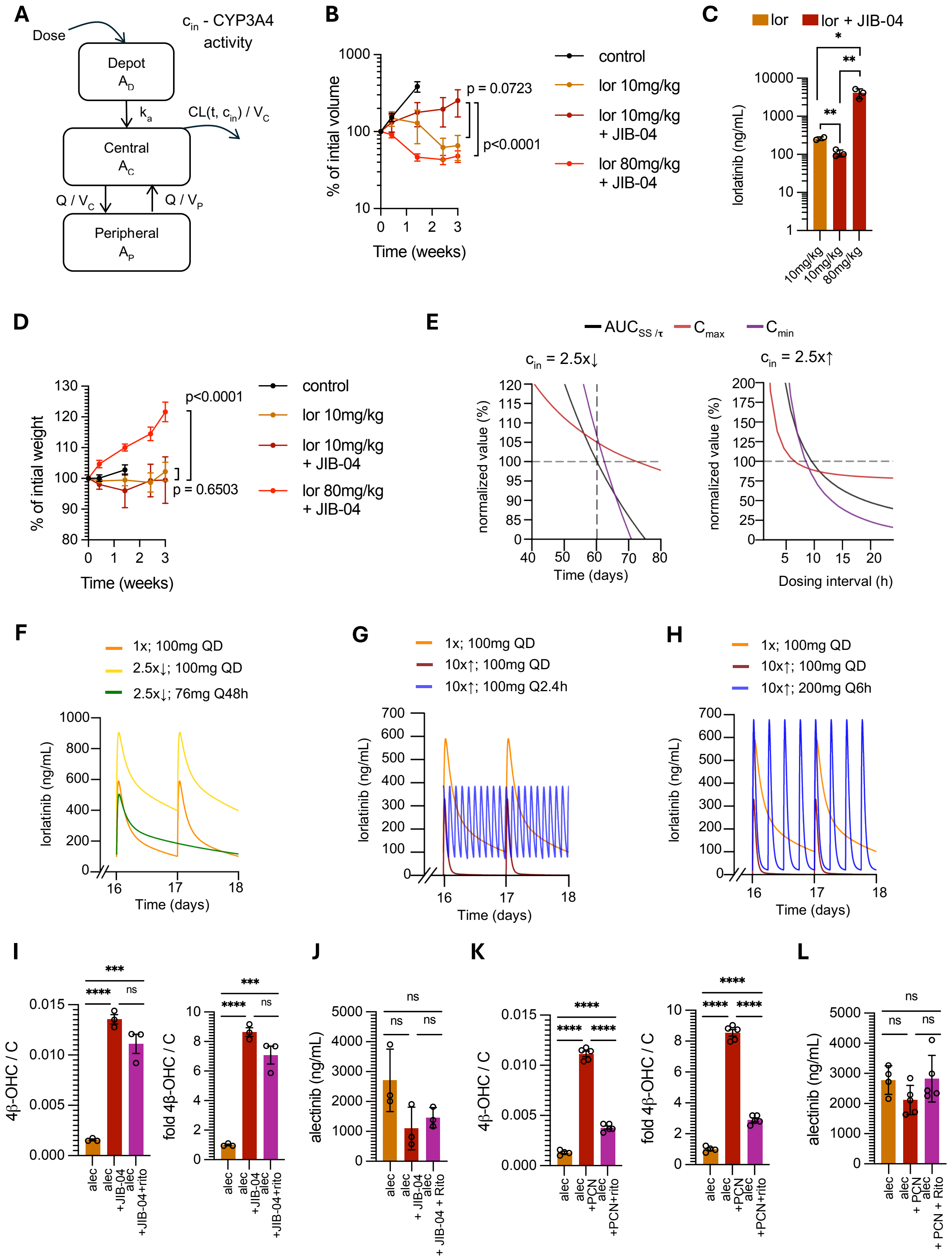

A
Dose
Depot A_D
k_a
Central A_C
Peripheral A_P
c_in - CYP3A4 activity
CL(t, c_in) / V_C
Q / V_C
Q / V_P
B
% of intial volume
Time (weeks)
control
lor 10mg/kg
lor 10mg/kg + JIB-04
lor 80mg/kg + JIB-04
p = 0.0723
p<0.0001
C
lor
lor + JIB-04
lorlatinib (ng/mL)
10mg/kg
10mg/kg
80mg/kg
D
% of intial weight
Time (weeks)
p<0.0001
p = 0.6503
control
lor 10mg/kg
lor 10mg/kg + JIB-04
lor 80mg/kg + JIB-04
E
AUC_SS/τ
C_max
C_min
c_in = 2.5x↓
c_in = 2.5x↑
normalized value (%)
Time (days)
Dosing interval (h)
F
1x; 100mg QD
2.5x↓; 100mg QD
2.5x↓; 76mg Q48h
lorlatinib (ng/mL)
Time (days)
G
1x; 100mg QD
10x↑; 100mg QD
10x↑; 100mg Q2.4h
lorlatinib (ng/mL)
Time (days)
H
1x; 100mg QD
10x↑; 100mg QD
10x↑; 200mg Q6h
lorlatinib (ng/mL)
Time (days)
I
4β-OHC / C
fold 4β-OHC / C
alec
alec +JIB-04
alec +JIB-04+rito
J
alectinib (ng/mL)
alec
alec + JIB-04
alec + JIB-04 + Rito
K
4β-OHC / C
fold 4β-OHC / C
alec
alec +PCN
alec +PCN+rito
L
alectinib (ng/mL)
alec
alec + PCN
alec + PCN + Rito

**Supplementary Figure 5: Mitigation strategies for altered CYP3A4/11 activity**

**A.** Schemata of mathematical model showing compartment model for PK of lorlatinib. **B.** Change in tumor volume of H3122 subcutaneous xenografts over time under indicated treatments. Y axis shows the % volume, normalized with initial volume measurements at the start of treatment. n = 8 tumors for control, n = 4 tumors for lor 10mg/kg and n = 6 tumors each for combination with JIB-04. p-value represents treatment term of 2-way ANOVA with Sidak's test. **C.** Plasma concentrations of lorlatinib from mice with indicated treatments for 3 weeks. n = 2 for lor 10mg/kg and n = 3 for lor combination with JIB-04. Unpaired t-test was performed. **D. %** change in weight of mice with indicated treatments relative to treatment start. n = 4 mice for control, n = 2 for lor 10mg/kg and n = 3 for combination. p-value represents treatment term of 2-way ANOVA with Sidak's test. **E.** Dynamics of PK metrics with change in dosing interval at indicated $c_{in}$. Values are normalized as percentage to that at $c_{in}$ of 1 with SOC dosage/interval. **F-H.** Concentration-time curves of lorlatinib at indicated $c_{in}$ values and dosage/interval (noted as $c_{in}$ ; dosing). **I.** Ratio and fold of 4β-OHC/C ratio where ratio in groups is normalized with ratio in alec alone group from mice treated with indicated drugs for 3.5 weeks. n = 3 mice for all groups. Unpaired t-test was performed. **J.** Alectinib concentrations in plasma after indicated treatments. n = 3 mice for all groups. Unpaired t-test was performed. **K.** Ratio and fold of 4β-OHC/C ratio where ratio in groups is normalized with ratio in alec alone group from mice treated with indicated drugs for 5.5 weeks. n = 4 mice for alec alone and n = 5 mice for rest. Unpaired t-test was performed. **L.** Alectinib concentrations in plasma after indicated treatments. n = 4 mice for alec alone and n = 5 mice for rest. Unpaired t-test was performed. Mean ± SEM are shown for all, *$p < 0.05$, **$p < 0.01$, ***$p < 0.001$, and ****$p < 0.0001$.
