## Supplementary Methods for "Pharmacokinetic Acceleration via CYP3A4 Hyperactivation as a Clinically Actionable Mechanism of Targeted Therapy Resistance in NSCLC"

Sex as a biological variable. For mouse studies, both male and female animals were included. While most experiments were performed in females, comparable findings were observed in both sexes.

qPCR. Following 48 hours treatment of Huh7 or AML12 cells with JIB-04 (1μM), RNA was extracted using RNeasy Mini kit (Qiagen). cDNA synthesis (SensiFAST cDNA synthesis kit, BioLine, Cat# BIO-65053) and qPCR (SensiFAST SYBR Hi-ROX Kit, BioLine, Cat# BIO-73005) were performed as per kit protocols. Primers used were: CYP3A4 (Forward - 5' GCC AAA GAA TCA ATT AGG CCC ATC T 3', Reverse – 5' TGA CTG TGC AAA ATA CTT CCC CA 3') and housekeeping control RPL19 (Forward – 5' GATCCGGAAGCTCATCAAAG 3', Reverse – 5' GGCTGTACCCTTCCGCTTAC 3') for the human hepatocyte cell line, Huh7; Cyp3a11 (Forward – 5' GAC AAA CAA GCA GGG ATG GAC 3', Reverse – 5' CCA AGC TGA TTG CTA GGA GCA 3') and housekeeping control b-actin (Forward – 5' CAC TGT CGA GTC GCG TCC A 3', Reverse – 5' GTC ATC CAT GGC GAA CTG GT 3') for the murine hepatocyte cell line, AML12.

Secreted luciferase assay (GLuc) for in vivo tumor growth monitoring. A few microliters of blood were collected from tail vein incision of experimental mice into heparin coated tubes, centrifuged at 2000g for 10 minutes, supernatant plasma was collected and stored at -80 until used for the assay. Plasma samples were diluted 1:100 in PBS and 20ul of this was loaded onto white opaque bottom 96 well plates. 100ul of 5ug/mL Coelenterazine (GoldBio CZ) was injected using the Promega GloMax luminometer and luminescence was read. PBS injected with substrate was used as blank. Luminescence was measured in blood collected weekly and % of initial luminescence was calculated by normalizing all subsequent luminescence with luminescence of the sample collected on the day of starting the drug treatments.

Bulk RNA sequencing. Bulk RNA sequencing was performed using mouse liver samples. Snap-frozen livers were crushed into powder using mortar and pestle on dry ice and pouring liquid nitrogen. Powder was dissolved in Trizol (Invitrogen, Cat# 15596025), followed by chloroform-ethanol precipitation to harvest RNA. Nanodrop and Agilent 2100 Bioanalyzer were used to assess the concentration and quality of RNA. RNA samples were submitted to Novogene for sequencing and analysis. Briefly, HiSat2 was used for alignment to the Mus musculus (GRCm39/mm39) genome and differential gene expression analysis was performed using EdgeR and DeSeq2. Of this, genes with $-1.5<\log 2$ fold change$<1.5$ and p-adjusted$<0.05$ were selected and z-scores were calculated from FPKM values. This gene list was used to create heatmaps using Morpheus and, KEGG and Reactome barplots using ShinyGO, for genes upregulated in JIB-04 group vs control. For the heatmap of metabolism-related genes, genes from KEGG pathway analysis by ShinyGO were used. Lastly, GSEA was performed, and a representative enrichment plot was shown.

BrDU histology. Tissue samples were fixed in 10% formalin for 24 hours and then embedded in paraffin. For BrDU labeling, 10mg/mL BrDU in PBS was intraperitoneally injected in mice. The mice were euthanized 30-45 minutes later. The processing, embedding and staining were performed as previously described[1].

Quantification of 4β-hydroxy cholesterol and cholesterol. Reagents and chemicals - LC-MS grade water, methanol and iso-propanol (IPA) are from VWR; Formic acid is from Fisher Scientific (Waltham, MA). Internal standards (IS) Cholesterol (2,3,4-13C3,99%) is purchased from Cambridge Isotope Lab (Tewksbury, MA) and 4β-hydroxy Cholesterol-D7 is purchased from Cayman Chemical (Ann Arbor, MI). Sample Preparation - All processes were carried out on ice.

An aliquot of IS mixture was added into each sample. For mouse plasma, 500µL precooled 100% IPA extraction solvent (kept in the -80°C freezer at least one hour prior to extraction) was added to 100 µL sample (diluted in PBS) for protein precipitation. The samples were vortexed and centrifuged at 18,800 × g (Microfuge 22R, Beckman Coulter) at 0°C for 10 minutes. Then, the samples were incubated for 30 minutes in a -80°C freezer to increase IPA extraction. After that, the supernatant was transferred to a new a microcentrifuge tube and dried in a speedvac. Dried compounds were eventually re-dissolved in 20µL methanol. For human plasma, protocol was adapted from Yaodong Xu et.al[2]. Briefly, 40µL 80% MeOH was added to 10µL aliquots, then followed by a hydrolysis with 1M KOH (95% MeOH and 5% $H_2O$)[1]. After addition of 150µL of 1M KOH, the samples were vortexed and incubated for 45 minutes in 37°C. Then 600µL hexane was added to the samples for extraction. Similar to mouse samples, after vortex, incubation and centrifugation, the supernatant was dried and re-dissolved in 20µL methanol. LC-MS - Liquid Chromatography-Mass Spectrometry (LC-MS) was performed using a UHPLC (Vanquish, Thermo Scientific) interfaced with a Q Exactive HF hybrid quadrupole-Orbitrap mass spectrometer (Thermo Scientific, San Jose, CA). Chromatographic separation was conducted on an Accucore C18 column (2.1 mm ID x 100 mm in length with 2.6 µm particle size) maintained at 40 °C. Separation was achieved using mobile phases A (100% H2O with 0.1% formic acid) and B (100% MeOH with 0.1% formic acid). The gradient was programmed as follows: 2 minutes at 50% B, then using a linear gradient of 50-80% B over 6 minutes, and then ramping up to 99% B after 7 minutes, washing with 99% B for 2 minutes, returning to 50% B over 0.1 minutes and re-equilbrating for 2.9 minutes for a total run time of 20 minutes. The injection volume was 5µL. The flow rate was set to 0.400 mL/min. For ion source, APCI (Atmospheric Pressure Chemical Ionization) was used for the ionization. The data analysis is performed using XCalibur 4.0.

*Quantification of drugs.* Plasma: Plasma samples for alectinib, M4, lorlatinib, sotorasib, osimertinib and brigatinib were stored at a temperature of −80°C until analysis. Samples underwent protein precipitation with pre-chilled acetonitrile containing internal standard erlotinib at 50 ng/ml. Samples were vortexed, centrifuged and 2 µl of supernatant fraction was injected on a Dionex-Ultimate™ UHPLC system coupled to a TSQ Quantiva triple quadrupole mass spectrometer (Thermo Scientific™, San Jose, CA) equipped with electrospray ionization. Analytes were separated on a reverse-phase C18 column over a 5.5-minute gradient. A ratio of drug area to internal standard was used to plot against linear regression calibration curve ranging from 1.0 ng/ml to 2,500 ng/mL to determine drug concentration utilizing Trace Finder (ver. 3.2) software. Tumor: 100 µg of the frozen tumor sample was homogenized in a bullet blender (NEXT ADVANCE, Troy, NY) utilizing stainless steel beads and 1 mL of water. Drug extraction was conducted by aliquoting 25 µL of tumor homogenate crashed with pre-chilled protein precipitation solution (acetonitrile containing internal standard erlotinib at 50 ng/ml). Samples were vortexed, centrifuged and 2 µL of supernatant was injected on a Dionex-Ultimate™ UHPLC system coupled to a TSQ Quantiva triple quadrupole mass spectrometer (Thermo Scientific™, San Jose, CA) equipped with electrospray ionization. A ratio of drug area to internal standard was used to plot against a linear regression calibration curve ranging from 10 ng/g to 25,000 ng/g to determine drug concentration utilizing Trace Finder (ver. 3.2) software.

*Math model.* We adopt the 2-compartment lorlatinib population pharmacokinetic (popPK) model presented in Chen *et. al.* and incorporate a novel personalization parameter, $c_{in}$, representing individual CYP3A4 activity[3]. The set of ODEs describing the compartment PK dynamics are as follows:

$$\frac{dA_D}{dt} = -k_a A_D$$

$$\frac{dA_C}{dt} = k_a A_D - CL(t, c_{in})A_C - \frac{Q}{V_C}A_C + \frac{Q}{V_P}A_P$$

$$\frac{dA_P}{dt} = \frac{Q}{V_C}A_C - \frac{Q}{V_P}A_P$$

Simulations were performed in Python. For all simulations, parameter values were set to population mean values reported in Chen *et. al.* (Supplementary Table 1). The $c_{in}$ parameter scales the clearance function as follows:

$$CL(t, c_{in}) = c_{in}[CLI + (CLMX - CLI)(1 - e^{-t/\tau})]$$

A value of $c_{in}$ = 1 models population average CYP3A4 activity and we increase or decrease $c_{in}$ to model higher or lower than average CYP3A4 activity respectively.

The baseline patient is defined as having $c_{in}$ = 1 and SOC dosing (100mg QD). PK metrics $C_{max,ss}$, $C_{min,ss}$, and $AUC_{ss}$ of the baseline patient were extracted at steady-state from the final 24-hour dosing interval and used as reference metrics.

Three independent strategies were evaluated to personalize dosing regimens with the goal of normalizing PK metrics to the reference metrics – adjusting dose, dosing interval, or both. The adjusted protocol was simulated to steady state, defined as the change in AUC 5 doses apart falling below 1%, and PK metrics were extracted.

To find the optimal dose adjustment, the dosing interval was held fixed at 24 hours. A SciPy gradient-based optimizer, L-BFGS-B, identified the optimal dose by minimizing the relative squared error between the patient's simulated steady-state $AUC_{ss}$ and the reference $AUC_{ss}$:

$$\min_{D} \left( \frac{AUC_{SS}(D, c_{in}) - AUC_{SS}^{ref}}{AUC_{SS}^{ref}} \right)^2$$

for doses in D ∈ [10, *max* (600, 200$c_{in}$)] mg and an initial guess of 100$c_{in}$. The upper bound scales linearly with $c_{in}$ since optimal dose ∝ $c_{in}$ in a linear PK system. This convexity guarantees a unique global minimum, so a single initial guess suffices.

To find the optimal dosing interval, the dose was held fixed at 100 mg. A SciPy bracketed root-finding algorithm, brentq, was used to find the optimal interval by solving: $\frac{AUC_{ss}}{\tau} - \frac{AUC_{ss}^{ref}}{\tau} = 0$ for interval lengths in $\tau \in [1,96]$ h. A unique root is guaranteed because $\frac{AUC_{ss}}{\tau}$ decreases monotonically with increasing $\tau$. The tolerance is set to 0.05 h, which guarantees convergence to a solution reported to the nearest 3 minutes.

Here, the optimization target was $\frac{AUC_{ss}}{\tau}$ rather than $AUC_{ss}$ because per-interval AUC scales proportionally with $\tau$, making it interval-dependent; whereas $\frac{AUC_{ss}}{\tau}$ normalizes by interval length and is comparable across different dosing intervals.

Finally, we simultaneously varied dose and dosing interval. The L-BFGS-B previously described was used to minimize the sum of equally weighted relative squared errors across all three steady-state PK metrics: $C_{max,ss}$, $C_{min,ss}$, and $\frac{AUC_{ss}}{\tau}$. The dose is again bounded to D ∈ [10, *max* (600, 200$c_{in}$)] mg but the interval is restricted to a tighter range $\tau \in [6, 48]$ h. To reduce sensitivity to local minima, the optimizer was run from a 3×3 grid of initial conditions: doses of [50 x $c_{in}$, 100 x $c_{in}$, 200 x $c_{in}$] mg crossed with intervals of [12, 24, 36] h, giving 9 starting points.
